# Oral L-arginine cures arginase 1-dependent chronic cutaneous leishmaniasis by redirecting the T helper cell response

**DOI:** 10.1101/2025.09.23.677696

**Authors:** Baplu Rai, David Barinberg, Chunguang Liang, Andrea Debus, Manfred Rauh, Sinan Kirici, Heidi Sebald, Myriam Grotz, Tobias Gold, Meik Kunz, Aline Bozec, Jochen Mattner, Sigrid Roberts, Ingeborg Becker, Christian Bogdan, Ulrike Schleicher

**Author notes:** shared senior and corresponding authorship (Christian Bogdan, Ulrike Schleicher). <u>Correspondence</u> Priv. Doz. Dr. Ulrike Schleicher and Prof. Dr. Christian Bogdan, Mikrobiologisches Institut, Universitätsklinikum Erlangen, Wasserturmstraße 3/5, D-91054 Erlangen, Germany.

## Abstract

*Leishmania (L.) mexicana*-induced cutaneous leishmaniasis (CL) is a neglected tropical disease characterized by localized chronic ulcers (LCL) and, in rare cases, by disseminated skin lesions (DCL). The therapeutic options for CL are currently limited, and the immune dysregulation leading to chronicity of disease is poorly understood. Here, we identified interleukin (IL)-10-dependent upregulation of arginase 1 (ARG1) in cutaneous CX3CR1^+^ myeloid cells as central immunometabolic determinant of chronic CL in *L. mexicana*-infected C57BL/6 wild-type (WT) mice. Deletion of *Arg1* in myeloid cells (*Arg1ΔCx3cr1*) enabled parasite control and clinical healing. Single-cell RNA sequencing revealed that ARG1, together with interferon-γ produced by T-helper 1 (Th1) cells, caused pathologic differentiation of Ly6C^high^ monocytes into inflammatory macrophages (iMACs) that simultaneously expressed ARG1, nitric oxide synthase type 2 (NOS2) and the chemokines CXCL9/10. These *Arg1^+^Nos2^+^Cxcl9/10^+^*iMACs induced a lasting depletion of L-arginine in the skin, served as parasite niche, and maintained a self-perpetuating cycle of host cell recruitment. In *Arg1ΔCx3cr1* mice, the ARG1^+^NOS2^+^ host cell niche for the parasite was diminished. Prophylactic or therapeutic oral L-arginine supplementation restored tissue arginine levels, reduced parasite burden, and prevented or resolved chronic disease. L-arginine-treated mice showed enhanced T cell expansion and Th1 differentiation, remained free of clinical relapses, and were resistant to reinfection. As skin lesion biopsies from *L. mexicana*-infected LCL and DCL patients demonstrated a similar pattern of Th1/Th2 cytokine, *Arg1* and *Nos*2 mRNA expression as seen in mice, we suggest metabolic reprogramming by oral L-arginine as a promising and easy-to-apply host-directed therapy for human *L. mexicana* CL.

**ONE SENTENCE SUMMARY:** Arginase 1-mediated arginine depletion accounts for chronic cutaneous leishmaniasis, which can be prevented and even cured by oral L-arginine therapy.

## INTRODUCTION

Cutaneous leishmaniasis (CL) is caused by different species of the protozoan parasite genus *Leishmania*, which are transmitted as extracellular, flagellated promastigotes by the bite of sand flies and transform into intracellular amastigotes after infection of mammalian hosts (*1*). The clinical course of infection ranges from harmless and self-healing, papular skin lesions at the site of infection to chronic and progressive ulcers that require prolonged local or systemic antiparasitic treatment (*2, 3*). The variable outcome of disease primarily depends on the infecting *Leishmania* species and the functional state of the immune system (*4*) and is governed by the entirety of cellular, humoral and physicochemical factors that form the local tissue microenvironment (“immunomicrotope”) (*5–7*).

Among the species causing CL, *Leishmania (L.) mexicana* can elicit chronic, long-lasting or even non-resolving localized skin lesions in experimental mouse models (*8–12*) and also in humans (*13*). In rare cases, *L. mexicana*-infected patients with a lack of cellular immunity to *Leishmania* antigens do not develop localized (LCL), but diffuse CL (DCL) characterized by disseminated, non-resolving cutaneous nodules that are difficult to treat (*2, 13–15*). Whereas C57BL/6 wild-type (WT) mice effectively cleared infections with most strains of *L. major* by a robust CD4+ Th1 immune response and interferon-γ (IFNγ) production (*4, 16*), C57BL/6 WT mice failed to resolve infections with *L. mexicana* (*9–12*). This has been linked to a weakened Th1 cell differentiation (*17*), enhanced IL-4/IL-13/STAT6 signaling (*18, 19*), and the expression of IL-10 (*20*). Further studies provided evidence that *L. mexicana* impaired the migration of monocyte-derived dendritic cells to the draining lymph node and the subsequent priming of Th1 cells for IFNγ production (*21*). Depletion of neutrophils, which were rapidly recruited to the site of *L. mexicana* infection, restored the Th1 response and prevented chronicity of infection (*12*).

Recent investigations have highlighted the crucial role of the immunometabolic microenvironment as a determinant of disease outcome in cancer (*22*), autoimmune diseases (*23*) or infections (*6, 24, 25*). One of the metabolic processes that is central to immune responses is the degradation of the semi-essential amino acid L-arginine (*26, 27*). Macrophages, the primary host cells harbouring *Leishmania* amastigotes (*5*), metabolize L-arginine predominantly via two competing enzymatic pathways with distinct functional outcomes: (a) type 2 or inducible nitric oxide synthase (NOS2 or iNOS) generates L-citrulline and nitric oxide (NO), which is critical for parasite killing and also exhibits immunoregulatory effects (*26*); (b) arginase 1 (ARG1) converts L-arginine into urea and ornithine, which is a precursor of polyamines and collagen that are necessary for cell proliferation and tissue repair (*27–29*). The balance between NOS2 and ARG1 expression within macrophages is primarily regulated by the local cytokine milieu: whereas Th1 cytokines (e.g., IFNγ, TNF) are strong inducers of NOS2, Th2 cytokines (e.g., IL-4, IL-13) as well as immunoregulatory cytokines (e.g., IL-10) predominantly enhance ARG1 (*5, 30–32*).

Considering the presence of a mixed Th1/Th2 cell-driven immune response with expression of IFNγ as well as IL-4/IL-10/IL-13 in mouse and human *L. mexicana* infections (*18, 19, 33, 34*), we hypothesized that Th2-driven ARG1 might play a pivotal role in causing chronicity of infection by creating a micromilieu that enables parasite persistence despite ongoing IFNγ and NOS2 expression. Our metabolic, single-cell RNA-sequencing (scRNAseq) and imaging analyses revealed that deletion of ARG1 restored L-arginine availability, prevented the self-amplifying generation of a ARG1^+^NOS2^+^ macrophage niche for the parasite at the site of infection, and allowed healing of otherwise chronic skin lesions. Furthermore, we observed that oral supplementation of C57BL/6 WT mice with L-arginine efficiently cured established chronic infections and resulted in long-lasting protective immunity against reinfection. Our study provides a comprehensive preclinical evaluation of oral L-arginine as a novel host-directed therapeutic strategy against chronic CL, bridging fundamental immunometabolic insights and translational application.

## RESULTS

### IL-10-induced ARG1 in myeloid cells accounts for chronicity of disease in *L. mexicana*-infected C57BL/6 mice

In a first set of experiments, we analyzed the mRNA and protein expression as well as function of the two isoenzymes of arginase, cytosolic ARG1 (primarily found in liver, myeloid and endothelial cells) and mitochondrial ARG2 (rather ubiquitously expressed, including myeloid and endothelial cells) (*28*), during the non-healing course of *L. mexicana* infection in C57BL/6 WT mice (**fig. S1A**). From day 20 to 40 post infection (dpi) onwards, ARG1 mRNA and protein significantly and continuously increased at the site of infection and correlated with the skin lesion development (**fig. S1A**), whereas the expression of ARG2 mRNA and protein remained low (**Fig. 1A-C**). Parallel to *Arg1*, we observed an mRNA upregulation of the ARG1-inducing cytokines *Il4*, *Il5*, *Il10* and *Il13* (**Fig. 1D**). A comparable, though less pronounced mRNA expression was seen in the draining lymph nodes (dLN) (**fig. S1B-D**). As time kinetics of *Il10* and *Arg1* expression were similar, we postulated that the previously reported healing of *L. mexicana* infections in IL-10-deficient mice (*20*) could be causally related to the absence of an ARG1 increase. Indeed, mice that lacked *II10* in CD4+ cells (*Il10ΔCd4*), spontaneously resolved an infection with *L. mexicana* (**fig.S1E-F**), released lower amounts of IL-10 upon restimulation *in vitro* (**fig.S1G**) and showed strongly reduced ARG1 mRNA and protein levels in the skin lesions, notably at later time points of the infection (**Fig. 1E and F**). Thus, IL-10 is one of the main stimuli upregulating ARG1.

**Fig. 1:**
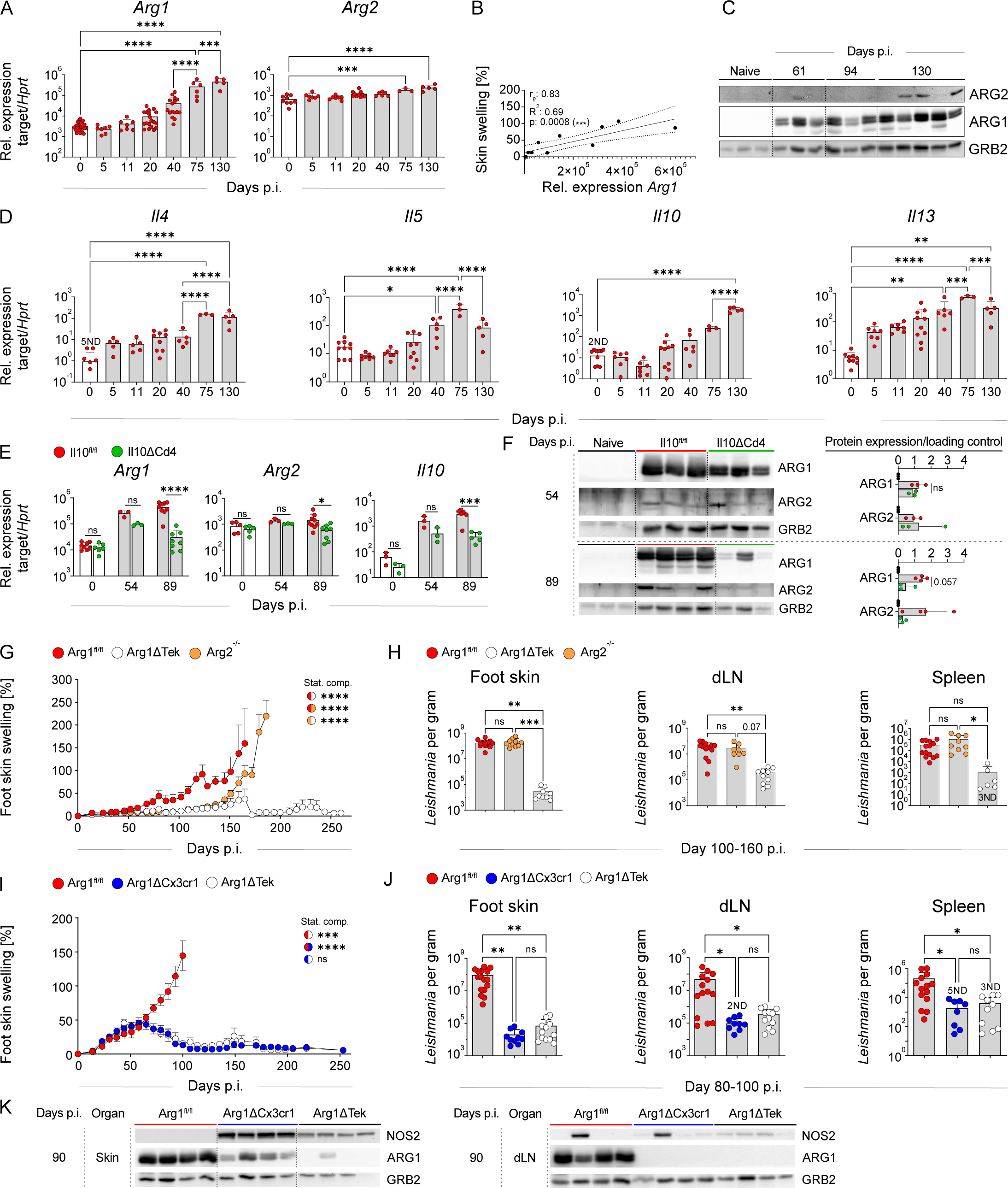
IL10-induced ARG1 in myeloid cells accounted for disease chronicity in *L. mexicana*-infected C57BL/6 mice. **(A, D, E)** mRNA expression in C57BL/6N WT **(A, D)**, *Il10ΔCd4* and *Il10^fl/fl^* **(E)** skin lesions at indicated days post infection (days p.i.) as determined by RT-qPCR. Bars show mean ± SD of 3-15 mice per time-point of 3 independent experiments. **(B)** Pearson correlation analysis of skin lesion size (skin swelling %) vs *Arg1* mRNA expression (Rel. expression *Arg1*) in the skin. 95% confidence band of the best fitting line is depicted. Each dot represents one mouse. **(C, F, K)** Representative Western blot analyses of ARG1, ARG2 and NOS2 in skin lesion lysates (80 μg protein/lane) of C57BL/6N WT **(C)**, *Il10ΔCd4* and *Il10^fl/fl^* **(F)**, *Arg1ΔTek*, *Arg1ΔCx3cr1* and *Arg1^fl/fl^* mice **(K)** at indicated days p.i. The protein expression was quantified by ImageJ software, with the ARG1 and ARG2 values being normalized to the corresponding loading control of GRB2 **(F**, right panel). Bars show mean ± SD of 3-4 mice per group. One of three independent experiments is shown. GRB2: growth factor receptor bound protein-2 as loading control. **(G, I)** Skin lesion development in *L. mexicana*-infected *Arg1ΔCx3cr1*, *Arg1ΔTek*, *Arg2^-/-^* and *Arg1^fl/fl^* mice. One of three independent experiments is shown. **(H, J)** Parasite numbers in the skin lesions, dLNs and spleens at indicated days p.i. were quantified by LD analysis in three independent experiments. Bars show mean ± SD of 9-16 mice per group. Statistical significance was determined using One-way ANOVA followed by Tukey’s multiple comparisons correction **(A, D, E, H, J)** and two tailed Mann-Whitney U test **(F)**. Comparisons between respective mouse groups were performed using a mixed-model two-way repeated measures ANOVA. Statistical significance between groups is illustrated using half-circles with the colours of the compared groups **(G, I)**. Significance levels are indicated by asterisks (*) according to the P value. ND, not determined; ns, not significant. p > 0.05; *p < 0.05; **p < 0.01; ***p < 0.001; ****p < 0.0001.

To assess the functional role of ARG1 and ARG2, C57BL/6 mice deficient for *Arg2* (*Arg2^-/-^*) (*35*) or lacking ARG1 in the endothelial and hematopoietic compartment (Arg1ΔTek) or the monocyte/macrophage compartment (*Arg1ΔCx3cr1*) (*32, 36–38*) were infected with *L. mexicana*. *Arg2^-/-^* mice developed progressive disease similar WT controls (*Arg1^fl/fl^*), although the onset of disease was delayed in *Arg2^-/-^* mice. In contrast, both conditional knockouts (KO) for *Arg1* (*Arg1ΔTek, Arg1ΔCx3cr1*) only showed mild skin swellings within the first 2 months p.i., which resolved thereafter along with a significant reduction of the tissue parasite load (**Fig. 1G-K**), of the numbers of neutrophils accumulating in the skin during the chronic phase (**fig. S1H and I**) and of the IL-4 protein concentrations in skin lysates at 90 dpi (**fig. S1J**). From these data we conclude that IL-10-induced ARG1 in CX3CR1+ myeloid cells drives chronic pathology and parasite survival in *L. mexicana* CL.

### ARG1 limits the production of NO by NOS2 in WT skin lesions

To elucidate the mechanisms, by which ARG1 drives pathology and non-healing disease, we focused on day 40 p.i., when lesion size and parasite load were not yet different between WT and *Arg1ΔCx3cr1* mice (**fig. S2A and B**). At this time-point, *Arg1* mRNA expression started to increase in WT skin lesions, which was paralleled by a significant upregulation of *Nos2* mRNA that was not seen in the infected skin of *Arg1ΔCx3cr1* mice (**Fig. 2A**). Using confocal laser scanning fluorescence microscopy (CLSFM), we found high levels of ARG1 and NOS2 protein in the cutaneous lesions of WT controls, whereas only a few, small clusters of NOS2 and ARG1 were detected in infected *Arg1ΔCx3cr1* mice (**Fig. 2B**). Considering that ARG1 and NOS2 were expressed in close proximity to each other in some regions of the WT skin lesions, we investigated whether ARG1 affected the *in-situ* production of NO as measured by direct (di-anthraquinone [DAQ] staining) or indirect detection methods (nitro-tyrosine staining) (*32*). As shown in **Fig. 2C and D**, similar amounts of NO were detected in WT and KO skin sections, despite the huge quantitative difference in the expression of NOS2 and ARG1 protein in both strains of mice. We conclude that in WT skin lesions high levels of ARG1 strongly limit the output of NO by the abundantly expressed NOS2, whereas in KO mice the near absence of ARG1 leaves the production of NO by the small amounts of NOS2 largely unaffected.

**Fig. 2:**
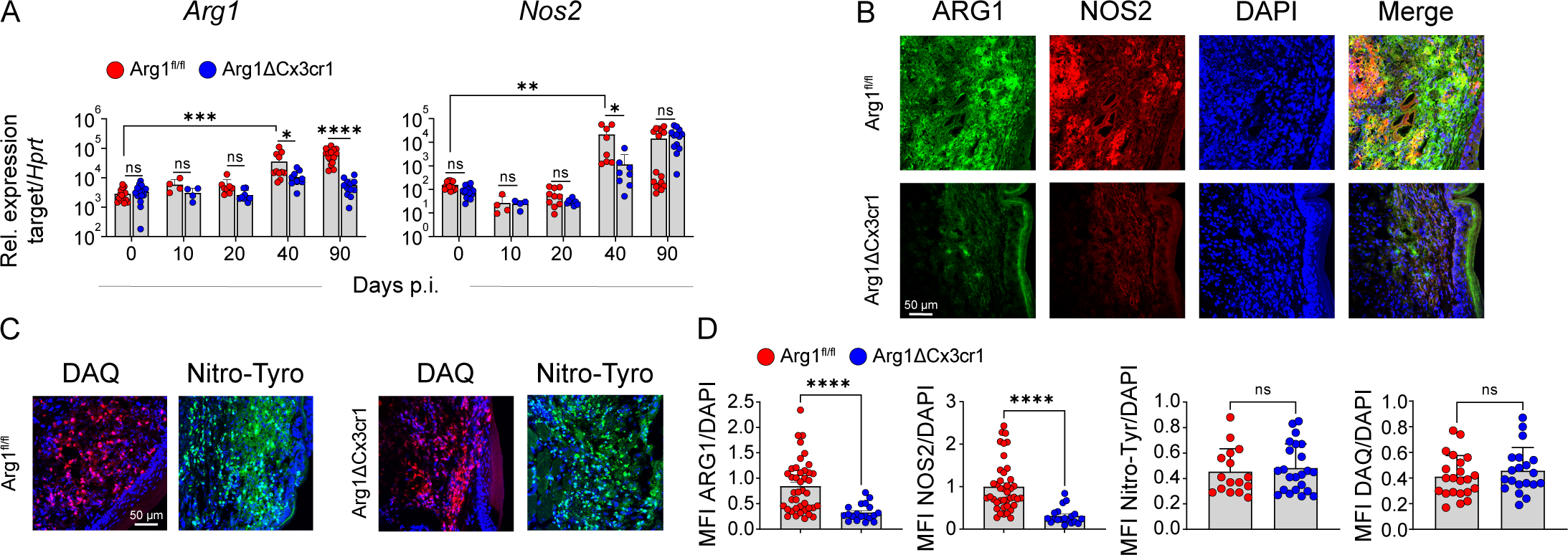
WT mice showed increased ARG1 and NOS2 levels, but the overall generation of NO was comparable in WT and *Arg1*-deficient mice at day 40 p.i. **(A)** Time course of *Arg1* and *Nos2* mRNA expression level in the skin lesions of *Arg1ΔCx3cr1* and WT (*Arg1^fl/fl^*) littermate controls at different days p.i. analyzed by RT-qPCR. Bars show mean ± SD of 4-15 mice per group from 4 independent experiments. **(B, C)** 5 μm serial sections of infected skin lesions (days 40 p.i.) were prepared and stained for ARG1 (green), NOS2 (red), DAQ (red), nitrotyrosine (green) and DAPI (Blue). Nitrotyrosine (Nitro-Tyro) and DAQ staining reflect NO levels *in situ*. DAQ and nitrotyrosine stainings are shown as merged images together with DAPI. Representative images (LSM700) from two independent experiments with 3-4 mice per group are shown. **(D)** ARG1, NOS2, nitrotyrosine, and DAQ stainings (days 40 p.i.) were quantified using Zeiss LSM700 Zen blue software and normalized to DAPI nuclear staining. Mean fluorescence intensity (MFI) was calculated from two independent experiments with 3-4 mice per group and each dot represents a single field of view of microscopic images. Statistical significance was determined using One-Way ANOVA followed by Tukey’s multiple comparisons correction **(A)** or two-tailed Mann-Whitney U test **(D)**. ns (not significant), p > 0.05; *p < 0.05; **p < 0.01; ***p < 0.001; ****p < 0.0001.

### ARG1 promotes differentiation of inflammatory monocytes into ARG1+NOS2+ inflammatory macrophages

To define the cell populations that express ARG1 and NOS2 and to study a potential impact of ARG1 on the differentiation of myeloid cells, we conducted single-cell RNA-sequencing (scRNA-seq) analysis of total viable cells from skin lesions of *Arg1^fl/fl^*WT and *Arg1ΔCx3cr1* mice on day 40 p.i. using the 10x Genomics platform. Visualization of scRNA-seq data of 2,067 WT and 1,694 KO cells using uniform manifold approximation and projection (UMAP) identified 7 cell clusters, of which the myeloid cell cluster was the largest (**Fig. 3A**). Sub-clustering of the myeloid compartment revealed 13 subpopulations (m0-m12) that differed in their transcriptomic profile; seven of them were strongly enriched in the infected WT skin (m0-m4, m7, m11) (**Fig. 3B and C**).

**Fig. 3:**
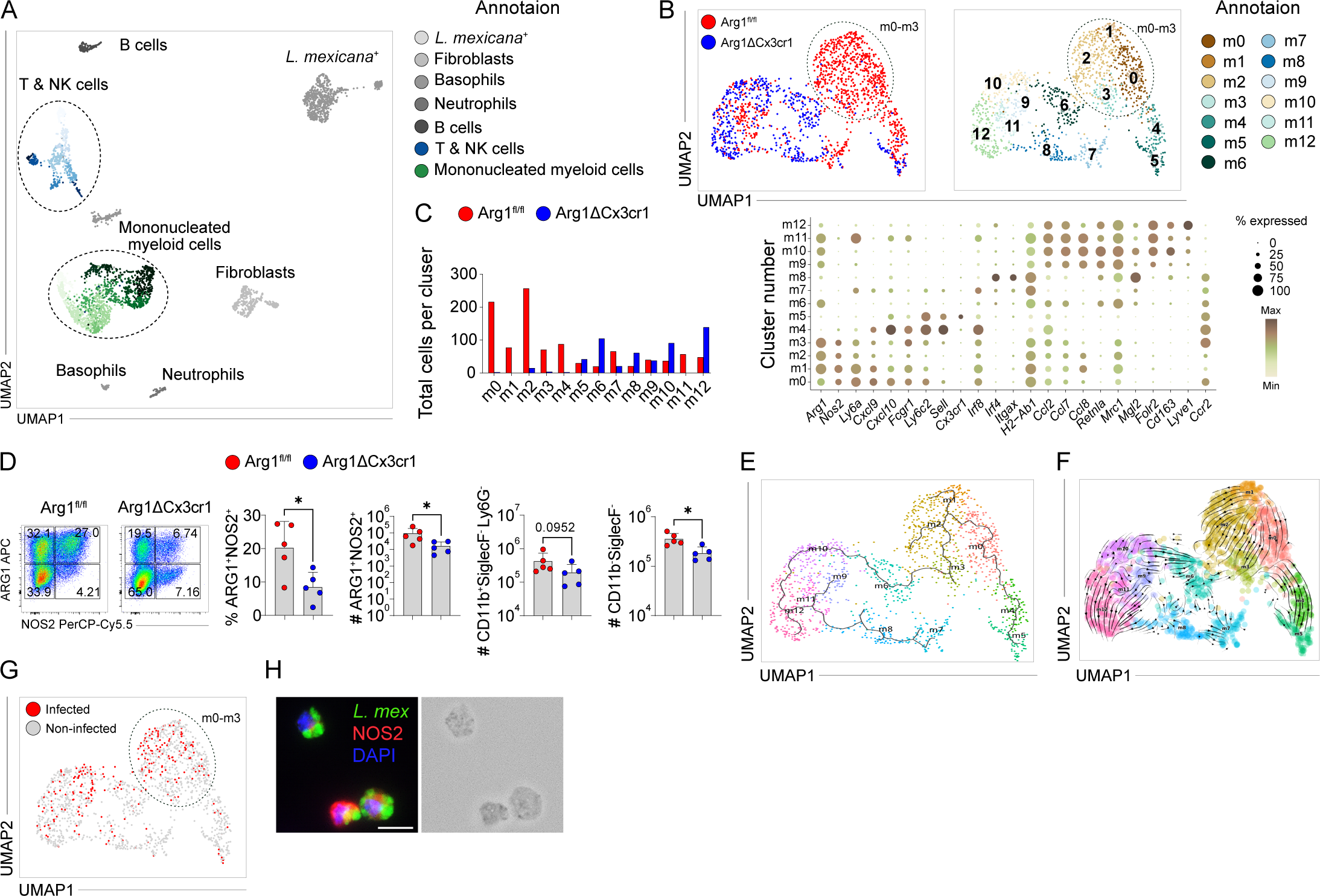
ARG1 promoted pathogenic differentiation of inflammatory monocytes towards ARG1^+^NOS2^+^ macrophages (iMAC) representing a cellular niche of *Leishmania*. **(A)** An overview UMAP was generated to visualize the distribution of all immune cell populations identified by scRNA-seq (10x Genomics) of viable cells pooled from skin lesions (3 mice per group) of *Arg1ΔCx3cr1* (KO) and *Arg1^fl/fl^* (WT) littermates at day 40 dpi. **(B)** UMAP representation of myeloid populations following sub-clustering into 13 distinct clusters, with the distribution of WT and KO cells visualized in red and blue, respectively. (**C**) The bar graph (left panel) depicts the number of cells in each myeloid subpopulation. The dot plot (right panel) illustrates the expression of key marker genes used to identify specific myeloid cell types. scRNA-seq data were processed using Cell Ranger (version 4.0) and analyzed with the R-based Seurat v4 pipeline. **(D)** Skin lesion samples were pooled from 3 - 4 mice per group in each experiment. Single-cell suspensions of skin cells were prepared, stained, and analyzed by flow cytometry. Flow cytometric analysis of ARG1 and NOS2 expression within CD11b⁺SiglecF⁻Ly6G⁻ populations was performed. The percentages and absolute numbers of ARG1⁺NOS2⁺, CD11b^+^SiglecF^-^Ly6G^-^ and CD11b⁻SiglecF⁻ populations were calculated using FlowJo software. Representative FACS plots and quantifications are shown from five independent experiments. **(E)** Pseudotime analysis of myeloid clusters using Monocle 3. **(F)** Trajectory analysis of myeloid clusters using RNA velocity. **(G)** UMAP showing *L. mexicana* positive cells. **(H)** Immunofluorescence staining of SiglecF^-^CD11b^+^ARG1^+^NOS2^+^ sorted cells stained for DAPI (Blue), *L. mexicana* (green), and NOS2 (Red). Merge (left) and bright field image (right). The images were taken with a Keyence fluorescence microscope (40× objective lens with 3× zoom function). Statistical significance was determined using two-tailed Mann-Whitney U test. ns (not significant), p > 0.05; *p < 0.05; **p < 0.01; ***p < 0.001; ****p < 0.0001.

Clusters m0-m3 represented 41.5% of all myeloid cells and were classified as **inflammatory monocyte-derived macrophages (iMAC)** based on their expression of *Ccr2*, *Ly6c2*, *Irf8*, *Ly6a (Sca1)*, *H2-Ab1 (MhcII)* and *Nos2*. Of note, these were the predominant *Nos2*+ populations in the skin (fig. S3A) and they all simultaneously expressed *Arg1* (**Fig. 3C**). Additionally, iMAC showed upregulation of many chemokines including *Cxcl2*, *Cxcl9* and *Cxcl10* as well as *Ccl2*, *Ccl5*, *Ccl7*, *Ccl8* and *Ccl24* (**Fig. 3C** and data not shown). When preparing single cell suspensions from skin lesions at 40-45 dpi, we noticed that the number of cells obtained from WT lesions was roughly 40-50% higher than in the case of *Arg1ΔCx3cr1* lesions (WT: 2.57×10^6^ ± 4.02×10^5^ vs KO: 1.45×10^6^ ± 1.86×10^5^, mean ± SEM, n=7). Flow cytometry revealed that an increased number of myeloid (CD11b^+^SiglecF^-^Ly6G^-^) and non-myeloid cells (CD11b^-^SiglecF^-^) contributed to the more prominent accumulation of cells in the skin lesion of WT mice (**Fig. 3D**, myeloid gating strategy, see **fig. S3C**). The percentage as well as the absolute number of ARG1+NOS2+ macrophages within the CD11b^+^SiglecF^-^Ly6G^-^ myeloid compartment was significantly increased in the skin of infected WT compared to *Arg1ΔCx3cr1* mice (**Fig. 3D**). RT-qPCR analysis of the infected skin tissue verified augmented expression of chemokines (*Ccl2*, *Cxcl9*, *Cxcl10*) in WT lesions at 40-45 dpi (**fig. S3B**). In addition to iMAC, prominent *Arg1* mRNA was also found in **M2-like macrophage (Mrc1^+^, Retnla^+^) clusters** m6 and m9-m11 (**Fig. 3C**), which, with the exception of m11, are likely to account for the residual ARG1 levels observed in the skin lesions of *Arg1ΔCx3cr1* mice (**Fig. 1K**). Clusters 7 and 8 were categorized as **migratory, conventional dendritic cells (cDC1 and cDC2)** based on their expression of *Ccr2*, *Itgax (Cd11c)*, high *H2-Ab1 (MhcII)* as well as *Irf8* or *Irf4* and *Mgl2 (Cd301b)* (*39–42*) (**Fig. 3C**). The *Cx3cr1^+^* populations m4 and m5 were identified as **Ly6c2^+^ inflammatory monocytes** recruited to the skin via *Sell (Cd62l)*.

Whereas the m5 population was equally present in WT and KO skin, cluster m4 showed a strong pro-inflammatory profile (high *Cxcl10*) and was almost absent in *Arg1ΔCx3cr1* skin lesions (**Fig. 3C**). This observation prompted us to investigate, whether recruited inflammatory monocytes (m5) under WT conditions (ARG1-sufficient microenvironment) undergo a specific differentiation process via m4-monocytes towards m0-m3 iMAC populations. To prove this differentiation trajectory, we applied two bioinformatic analyses. Pseudo time analysis indicated the differentiation from m5 to m4 and further to m0-m3 (**Fig. 3E**), whereas RNA velocity analysis provided the directionality of this differentiation process (**Fig. 3F**). Thus, we concluded that ARG1 boosts the differentiation of CX3CR1^+^ inflammatory monocytes, which were recruited to the site of *L. mexicana* infection, into ARG1^+^NOS2^+^chemokine^+^ iMAC.

### ARG1^+^NOS2^+^ inflammatory macrophages function as host cell niche for *L. Mexicana*

As ARG1 restricts the activity of NOS2 and thereby the production of leishmanicidal NO (**Fig. 2**) (*5*), we hypothesized that ARG1^+^NOS2^+^ iMAC promote disease progression by acting as a replicative niche for *Leishmania* rather than control the infection by killing the parasite. Indeed, when analyzing which of the myeloid clusters m0-m12 of our scRNAseq analysis were positive for *Leishmania* transcripts, parasite genes were selectively enriched in *Arg1*^+^ clusters, including m0-m3, m6, and m9-m11 (**Fig. 3G**). Furthermore, on day 40 p.i., sorted CD11b^+^ SiglecF^-^ARG1^+^NOS2^+^ cells from the skin of infected WT mice were fully packed with *L. mexicana* amastigotes as shown by immunofluorescence microscopy (**Fig. 3H**; see **fig. S3D** for the gating strategy of sorted ARG1^+^NOS2^+^ cells). Thus, the huge number of iMAC in WT skin lesions serve as host cells for *Leishmania* parasites.

### IFNγ drives the development of ARG1^+^NOS2^+^ inflammatory macrophages

Gene set enrichment analysis of clusters m0-m3 indicated that iMAC may also act as antigen-presenting cells towards T cells and respond to IFNγ (**fig. S2C**), which prompted us to take a closer look at the T/NK cluster of the scRNA-seq data (**Fig. 3A**). Sub-clustering revealed eight distinct subpopulations, with five of them (0, 2–5) being clearly enriched in infected WT skin lesions (**Fig. 4A**). These included CD4^+^ *Rora*^+^ T cells (0), *Ifng*^high^ Th1 cells (2), NK cells (3), Ly6c2^+^ γδ T cells (4) and CD8^+^ T cells (5). Cluster 1 (*Il17a^+^* γδ T cells) and cluster 6 (*Il4^+^Il5^+^Il10^+^Il13^+^*Th2 cells) were comparably present in WT and KO skin, whereas cluster 7 (representing ILC2) was more prevalent in the KO skin. Hence, the T/NK compartment showed a strong *Ifng* expression in the WT skin with a prominent accumulation of *Ifng*^+^ Th1 cells, which was not observed in *Arg1ΔCx3cr1* skin lesions. A transient increase in the mRNA expression of the Th1 cytokines *Ifng* and *Tnf* in WT skin lesions was also detected by RT-qPCR (**fig. S4A**). Using flow cytometry, we also found a higher percentage and absolute number of CD4^+^ T cells and a higher frequency of IFNγ-producing CD4^+^ T and NK cells in infected WT as compared to KO skin lesions on day 40 p.i. (**Fig. 4B**; T cell gating strategy see **fig. S4B**). Consistent with this observation, the IFNγ protein concentrations in skin lesion lysates were elevated in *L. mexicana*-infected WT mice compared to *Arg1ΔCx3c1* mice on day 40 p.i. (**Fig. 4C**).

**Fig. 4:**
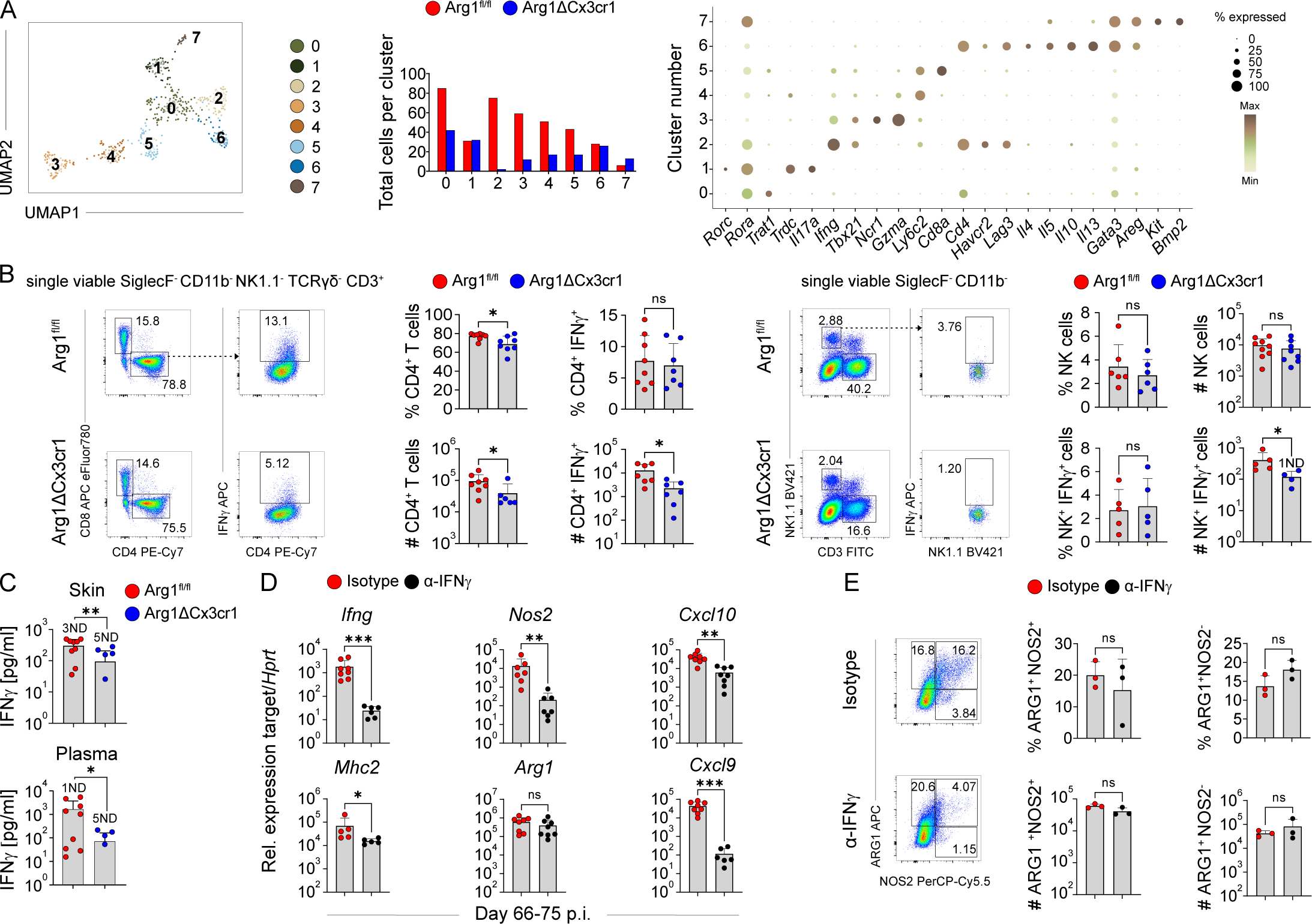
Increased IFN-ψ production by CD4⁺ T cells in the skin promoted the generation of ARG1⁺NOS2⁺ iMAC. **(A)** Subclustering of the lymphoid (T/NK) cluster and UMAP representation of lymphoid subpopulations at 40 dpi (left panel). The bar graph shows the number of cells in each lymphoid subpopulation, with WT and KO cells represented in red and blue, respectively (middle panel). The dot plot presents common markers used to identify immune cell types (right panel). scRNA-seq data were analyzed using Cell Ranger (version 4.0) and the R-based Seurat v4 scRNA-seq analysis pipeline. **(B)** Flow cytometry of T cell populations (SiglecF⁻CD11b⁻NK1.1⁻TCRγδ⁻CD3⁺) was performed, with further classification into CD4⁺ subsets. CD4^+^ T cells were subsequently analyzed for IFN-γ production (left panel). Flow cytometric analysis of NK cell populations (SiglecF⁻CD11b⁻CD3⁻NK1.1⁺), further assessed for IFN-γ production (right panel). Percentages and absolute numbers of respective populations were calculated using FlowJo software. Representative FACS plots and quantification from 5 - 8 independent experiments are shown. Skin lesion samples were pooled from 3 - 4 mice per group at day 40 days p.i. **(C)** IFN-γ level was measured in skin lysates and plasma samples at 40 days p.i. using ELISA. **(D)** *Arg1*, *Nos2*, *Ifng*, *Cxcl9*, Cxcl10, and *Mhc2* mRNA expression in the skin lesions of anti-IFN-γ-or isotype-treated C57BL/6N WT mice at 66-75 days p.i., measured by RT-qPCR. Bars represent mean ± SD of 5 - 8 mice per group. **(E)** Flow cytometry plots showing ARG1 and NOS2 expression within SiglecF⁻Ly6G⁻CD11b⁺ populations from mice treated with anti-IFN-γ antibody or isotype control. Percentages and absolute numbers were calculated using FlowJo software. Flow cytometry plots and quantification are from one independent infection experiment with three different time points of analysis (66–87 dpi). Skin lesion samples were pooled from 3 - 4 mice per group. **(B-E)** Statistical significance was determined using the two-tailed Mann-Whitney U test. ND, not determined; ns, not significant. Significance thresholds: p > 0.05 (ns); *p < 0.05; **p < 0.01; ***p < 0.001; ****p < 0.0001.

IFNγ was critical for monocyte-to-macrophage transition in an autoimmune disease model (*43*). Gene-set enrichment analysis of our *Ifng*^+^ Th1 cluster suggested an involvement of highly proliferative T cells in myeloid differentiation (**fig. S4C**). Therefore, we investigated whether differentiation of m5-monocytes towards m0-m3 iMAC was linked to the upregulation of IFNγ mRNA and protein in WT skin lesions around 40 dpi (**fig. S4A**). To this end, *L. mexicana*-infected C57BL/6 mice were treated with a suboptimal dose (50 µg per mouse) of a monoclonal neutralizing rat anti-mouse IFNγ antibody or the respective isotype control from 38 dpi onwards twice a week. Both groups of mice developed a similar course of infection (**fig. S4D**). Anti-IFNγ treatment caused a significant reduction of the mRNA levels of *Nos2* and of the chemokines *Cxcl9* and *Cxcl10*, which are mainly expressed by m4-monocytes and m0-m3 iMAC, in skin lesions at 66-75 dpi (**Fig. 4D**). Flow cytometry revealed a tentatively reduced percentage and absolute number of ARG1^+^NOS2^+^ iMAC within the CD11b^+^SiglecF^-^Ly6G^-^ myeloid compartment in the skin lesions of anti-IFNγ-treated mice, whereas percentage and number of ARG1^+^NOS2^-^ cells remained unaltered (**Fig. 4E**). Interestingly, anti-IFNγ treatment also strongly reduced the expression of *Ifng* mRNA (**Fig. 4D**), which might reflect a reduced recruitment and restimulation of Th1 cells due to the lower number of antigen-presenting MHCII^+^ARG1^+^NOS2^+^ chemokine^+^ iMAC.

Together, these data suggest a self-perpetuating cycle of inflammation, in which (1) ARG1 expression by *Cx3cr1^+^*recruited monocytes (m5) favors the differentiation into highly inflammatory *Cxcl10*^+^ m4-monocytes that subsequently attract T cells and more monocytes to the lesion; (2) recruited T cells produce IFNγ that promotes differentiation of m4-monocytes into m0-m3 *Arg1^+^Nos2^+^Cxcl9/10^+^MhcII^+^*iMAC; (3) chemokine-expressing iMAC further stimulate the infiltration of the skin lesions by T cells and monocytes and closely interact with T cells and activate them via MHCII-dependent antigen presentation. At the same time, iMAC serve as a habitat for *L. mexicana*.

### ARG1 activity changes the skin amino acid micromilieu

The prominent accumulation of ARG1^+^NOS2^+^ iMAC in the *L. mexicana*-infected skin lesions of WT mice leads to a high demand for L-arginine. To investigate whether changes in the amino acid metabolism contribute to the outcome of infection in non-healing WT versus healing Arg1ΔCx3cr1 mice, we determined the concentration of amino acids and polyamines in skin lesion lysates of both mouse strains at three different time-points p.i. using LC-MS. The comparative principal component analysis of 26 metabolites (19 amino acids, 3 polyamines, 4 others) showed no differences between WT and KO skin before and 35 days after infection. However, significant changes were observed on day 90 p.i. (**Fig. 5A**). In WT mice, induction of ARG1 led to the expected continuous rise of polyamines (e.g., putrescine, spermidine), whereas in Arg1ΔCx3cr1 mice, after an initial increase, the concentrations of polyamines dropped to pre-infection levels (**Fig. 5B**). Most of the amino acids measured showed a similar pattern as polyamines in WT and KO skin (**fig. S5A**), some others did not change significantly at all (data not shown). In contrast, the L-arginine concentration in the skin developed in a reverse manner: From day 0 to day 35, it dropped from 156.5±53.6 / 209.4±20.4 µmol/L to 18.1±4.9 / 28.4±3.2 µmol/L (WT/KO, mean ± SD). L-arginine remained at this low concentration on day 90 p.i. in WT skin lesions, but returned to naïve levels in healing *Arg1ΔCx3cr1* mice (176.3 ± 65.9 µmol/L) (**Fig. 5B**). In dLNs, the initial drop of L-arginine at 35 dpi was not seen, but at 90 dpi, the L-arginine concentration was strongly reduced in WT and normal in KO mice (**fig. S5B**). These data demonstrate that in *L. mexicana*-infected WT mice chronicity of disease correlated with a depletion of L-arginine in the infected tissues, whereas in self-curing *Arg1ΔCx3cr1* mice L-arginine levels normalized.

**Fig. 5:**
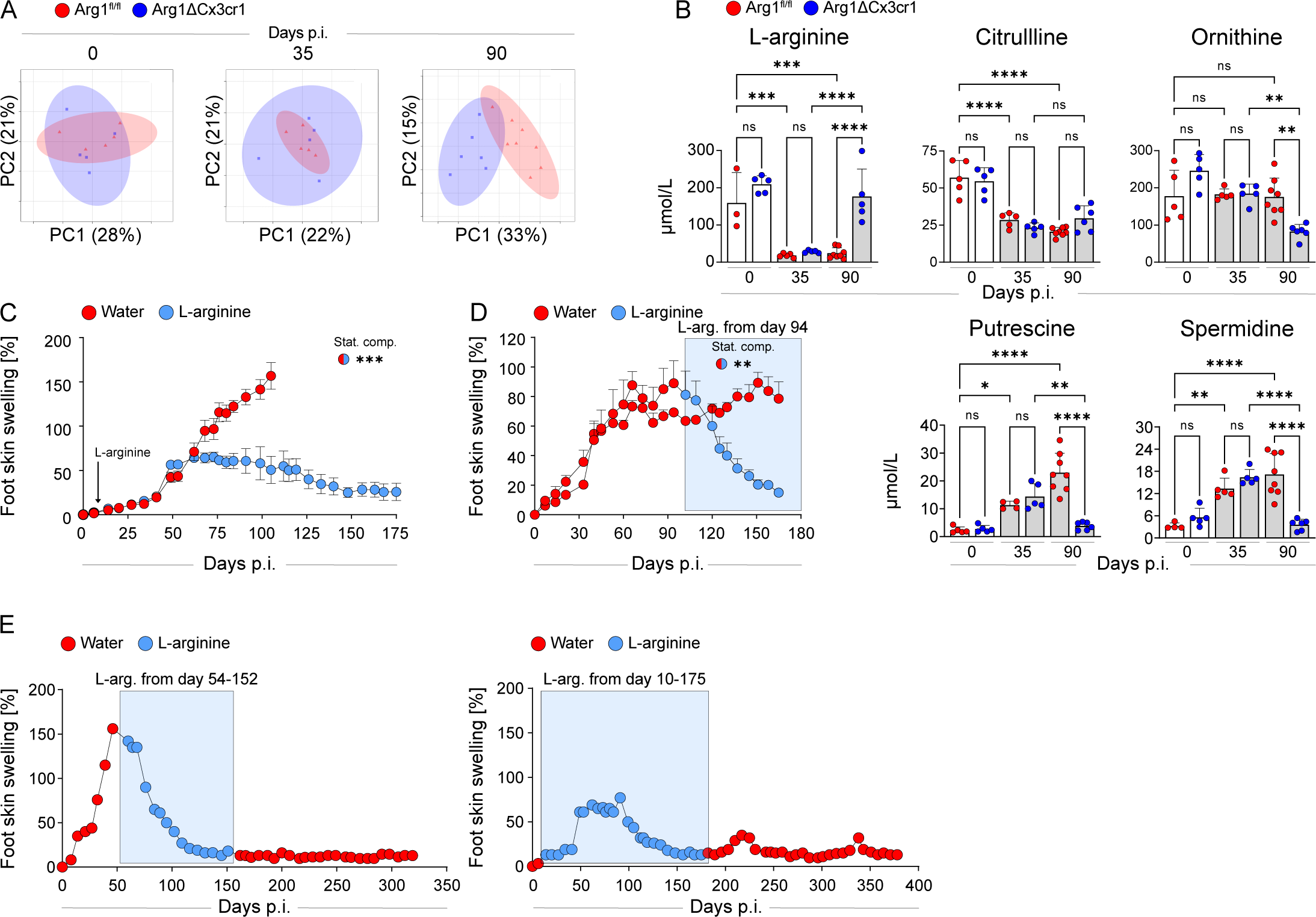
Resolution of chronic *L. mexicana* skin pathology by oral L-arginine supplementation. **(A)** Skin lysates from *Arg1ΔCx3cr1* (KO) and *Arg1^fl/fl^* (WT) littermate control mice were collected at different days p.i. and analyzed by LC-MS (5 - 8 mice / group). Downstream analysis of processed metabolomic data was performed using MetaboAnalyst 5.0, and principal component analysis (PCA) plots were generated based on 26 metabolites. **(B)** Quantification of metabolite levels at the infection site in *Arg1ΔCx3cr1* and *Arg1^fl/fl^* mice at different time points. Data are presented as mean ± SD from 3-8 mice per group. **(C)** Skin lesion development in *L. mexicana*-infected C57BL/6N mice treated with L-arginine (40 g/L in drinking water) starting on day 10 p.i. compared to untreated controls (water alone). Data shown represent one of two independent experiments (9 - 12 mice / group). **(D)** Skin lesion development of *L. mexicana*-infected C57BL/6N mice treated with L-arginine (40 g/L) versus water control. Treatment was initiated after mice had reached 100% increase of skin swelling. Data are representative of four independent experiments. **(E)** Long-term clinical course of infection after therapeutic (left panel) versus prophylactic application of L-arginine. For each regimen, a total of 3-4 mice, taken from independent infection experiments, were analyzed and monitored individually for 4-6 months. Shown are the skin lesion developments of one representative, individual mouse for each of the two treatment schemes. **(B)** Statistical analyses were performed using One-way ANOVA followed by Tukey’s multiple comparisons test. **(C, D)** Comparisons between respective mouse groups were performed using a mixed-model two-way repeated measures ANOVA. Statistical significance between groups is illustrated using half-circles with the colours of the compared groups. **(B, C, D)** Significance levels are indicated by asterisks (*) according to the *P* value. ns, not significant; *p > 0.05; *p* < 0.05; **p < 0.01; ***p < 0.001; ****p < 0.0001.

### Clinical resolution of chronic CL by oral L-arginine supplementation

Considering the correlation between L-arginine availability and the course of infection, we decided to test, whether oral supplementation of L-arginine via the drinking water (*23, 44*) is sufficient to prevent or revert chronic CL in WT mice. In a first experiment, *L. mexicana*-infected C57BL/6 mice received normal or L-arginine-containing drinking water from day 10 p.i. onwards. Both groups of mice had similar food and water uptake during the whole infection period (**fig. S6A**). Mice on normal water showed the expected non-healing disease, whereas 10 out of 12 L-arginine-treated mice developed only mild symptoms and clinically resolved the infection (**Fig. 5C**) along with a significant reduction of the parasite load (**fig. S6B**). Following oral L-arginine supplementation, the L-arginine concentrations in the healing skin lesions (93.3±47.5 µmol/L, mean ± SD, n=3), dLNs (29.7±2.5 µmol/L, mean ± SD, n=3) and plasma (172.4±40.5 µmol/L, mean ± SD, n=3) reached the level of uninfected naïve mice (skin: 152.3±21.8 µmol/L; dLN: 20.8±0 µmol/L; plasma: 144.9±25.8 µmol/L; mean ± SD, n=2-5) (**fig. S6C**). The concentrations of ornithine and citrulline were similar in the normal water and the L-arginine-treated group. Notably, application of L-arginine did not lead to the accumulation, but to significantly reduced levels of polyamines (putrescine, spermidine and spermine) (**fig. S6D**). Next, we explored, whether application of L-arginine can also cure established CL. When L-arginine was added to the drinking water of *L. mexicana*-infected mice with full-blown skin lesions, we observed a reduction of lesion sizes within 1 to 3 weeks of treatment and complete resolution of clinical disease in most of the mice (**Fig. 5D**; **fig. S7A**), which again was accompanied by a marked decrease of the tissue parasite burden (**fig. S7B**). Within 10 days of treatment, L-arginine levels normalized in the plasma. The concentrations of polyamines did not increase in L-arginine-treated mice, indicating that dietary L-arginine did not feed the arginase pathway, which is highly active at the beginning of the treatment (**fig. S7C**). In total, 30 out of 35 C57BL/6 mice from four independent experiments (i.e., 85.7%) showed clinical healing after therapeutic L-arginine application, whereas five mice did not respond for unknown reasons.

In seven mice that were successfully treated with L-arginine using the prophylactic (n = 3) or the therapeutic scheme (n = 4), L-arginine supplementation was stopped after resolution of clinical disease and the mice were monitored for additional 28 to 55 days (n = 3) or even 150 to 205 days (n = 4). While four of these mice showed mild transient skin swellings, there was no spontaneous relapse of disease in L-arginine treated and healed mice (**Fig. 5E**). *L. mexicana*-infected C57BL/6 mice that were cured by L-arginine treatment, were also protected from clinical disease after reinfection with *L. mexicana* (**fig. S7D**).

Finally, we investigated the impact of therapeutic L-arginine application on gene expression. We analyzed skin tissue samples from L-arginine-treated vs. untreated mice (n = 7 per group) on day 10-15 after initiation of L-arginine supplementation, when skin swellings were already reduced by 20 to 30% compared to untreated controls (**fig. S8A**). Using RT-qPCR, we found significantly increased mRNA copy numbers of *Il2* and Th1 associated genes including the transcription factor *Tbx21*, the cytokines *Ifnγ*, *Tnf*, and *Il12p35* as well as the effector molecule *Nos2* in the L-arginine-treated skin. There were no alterations in *Gata3*, *Il4*, *Il10*, *Foxp3*, and *Rorc* expression, and a trend towards decreased expression of *Il5, Il13,* and *Arg1* in skin lesions of L-arginine-treated mice (**Fig. 6A**). A similar pattern of mRNA expression was observed in dLNs, although induction of Th1 cytokines after L-arginine supplementation was weaker as compared to the skin site (**fig. S8B**). A bead-based multiplex ELISA-assay (LEGENDplex^TM^) comparing skin lysates of L-arginine-treated mice versus water controls confirmed the trend towards higher IL-2 and Th1 cytokine levels in response to oral L-arginine on protein level (**Fig. 6B**). The upregulation of Th1 cytokines in the L-arginine-treated skin was accompanied by a higher mRNA and protein expression of Th1-/IFNγ-induced chemokines such as CCL5 (RANTES), CCL11 (Eotaxin-1), CXCL9 and CXCL10, whereas other chemokines were not affected (**Fig. 6C** and **D**; data not shown). The upregulated chemokines are known to attract immune cells, e.g., T cells. We therefore analyzed the cellular composition of the infected skin of L-arginine-treated versus untreated mice. The number of viable cells retrieved from the lesions of L-arginine-supplemented mice was increased compared to control mice in each experiment, although the skin swelling of L-arginine-treated mice was already reduced (**Fig. 6E**, **fig. S8A**). Flow cytometry revealed that CD3^+^CD4^+^ and CD3^+^CD8^+^T cell numbers expanded after L-arginine treatment (**Fig. 6F**). Performing intracellular IFNγ staining, we detected a tentative increase of the percentage and absolute number of IFNγ^+^ cells within the cutaneous CD4^+^ and CD8^+^ T cell compartment after oral L-arginine application (**Fig. 6G**). Together, these data demonstrate that oral L-arginine supplementation can cure chronic CL in mice. Restoration of L-arginine levels in the infected skin supports the recruitment of T cells and shifts the CD4^+^ T helper response towards a protective Th1 phenotype.

**Fig. 6:**
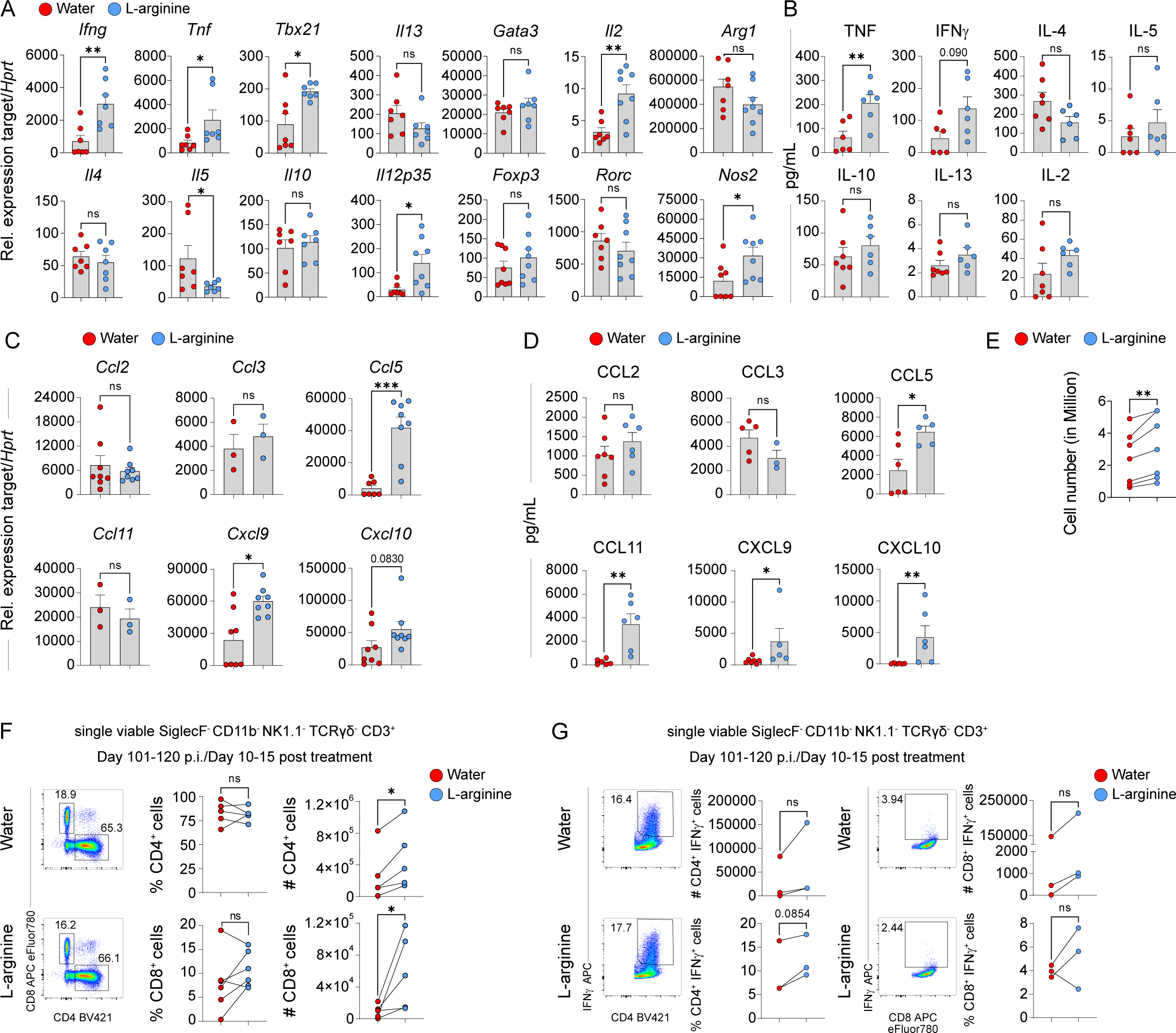
L-arginine-mediated resolution of disease was accompanied by a robust T cell expansion and a shift towards Th1. **(A, C)** mRNA expression (assessed by RT-qPCR) in skin lesions of *L. mexicana*-infected mice at day 10-15 of L-arginine treatment, which was started when the increase of skin lesion swelling had reached 100%. Data are presented as mean ± SEM (3-8 mice per group). **(B, D)** Cytokines and chemokine levels were quantified in skin lesion lysates by Legendplex analysis. Data are presented as mean ± SEM (3-8 mice per group). **(E)** *L. mexicana*-infected C57BL/6N WT mice with chronic CL lesions (100% increase of skin swelling) were split into two groups that either received L-arginine-containing drinking water or remained on normal water (control). Analyses were performed at day 10-15 after initiation of L-arginine treatment. Viable cells were isolated from skin lesions and pooled from 3-4 *L. mexicana*-infected mice per group and experiment. Depicted results are from 6 independent experiments. **(F, G)** Flow cytometric analysis of T cell populations from skin lesions (gated as SiglecF⁻CD11b⁻NK1.1⁻TCRγδ⁻CD3⁺ cells), further classified into CD4⁺ and CD8⁺ subsets. CD4⁺ and CD8^+^ T cells were analyzed for IFN-γ expression. Percentages and absolute numbers were quantified using FlowJo software. Representative flow cytometry plots and quantifications are shown from 3-5 independent experiments; skin lesion samples were pooled from 3 - 4 mice per group. Statistical analyses were performed using a two-tailed Mann-Whitney U test (A-D), parametric paired t test (E-G). ns, not significant; *p > 0.05; *p* < 0.05; **p < 0.01; ***p < 0.001; ****p < 0.0001.

### Upregulation of *Arg1* mRNA in lesions of *L. mexicana*-infected patients

The efficacy of L-arginine treatment in the mouse model suggests that dietary L-arginine might be a new therapeutic option for *L. mexicana*-infected CL patients. In a first translational approach, we measured mRNA expression of Th1/Th2 cytokines, *Arg1* and *Nos2* in skin lesions from Mexican patients with localized (LCL) or diffuse cutaneous leishmaniasis (DCL) as well as in healthy skin from uninfected Mexican volunteers living in the same area as the CL patients. Remarkably, we observed a similar expression pattern as seen in *L. mexicana*-infected skin lesions of mice. Both Th1 and Th2 cytokines as well as *Arg1* and *Nos2* were upregulated in the human lesions. In accordance with previous reports (*34, 45*), *Ifng* mRNA levels were higher in LCL as compared to DCL patients, but not absent in the latter (**Fig. 7**). These data suggest that *L. mexicana*-infected LCL and DCL patients might be responsive to an (adjuvant) treatment with L-arginine.

**Fig. 7:**
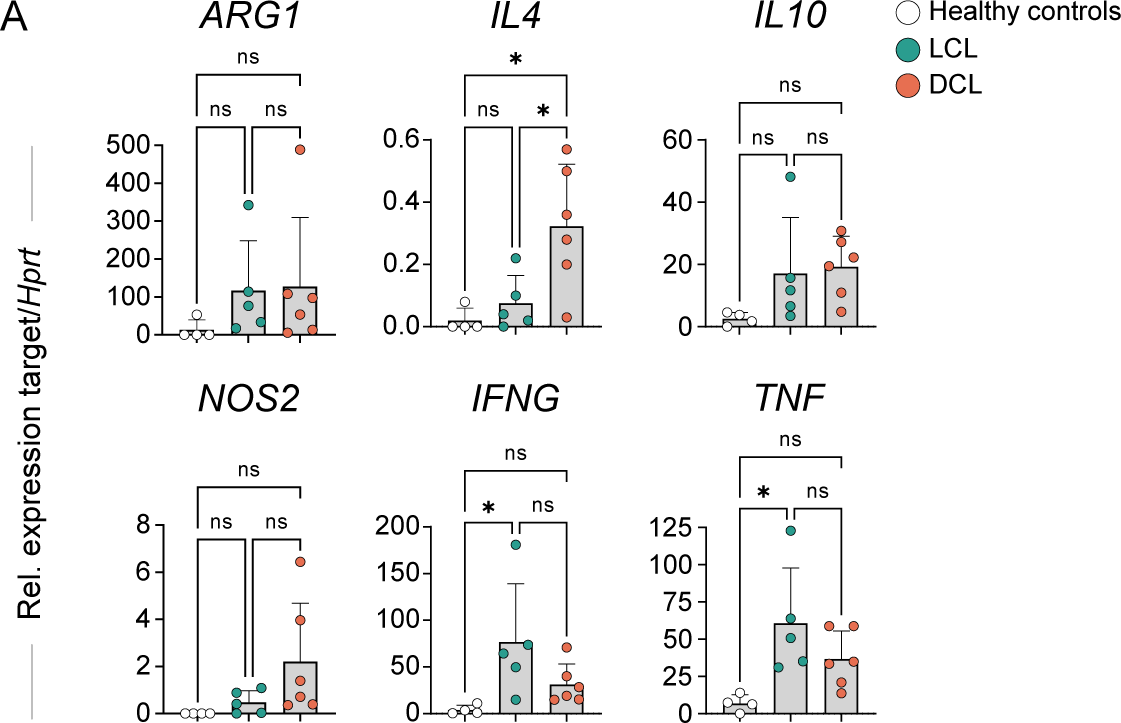
Elevated expression of *ARG1* and *NOS2*, along with concurrent induction of Th1/Th2 cytokines, were detected in human lesions of chronic CL. mRNA expression levels of *ARG1*, *NOS2*, *IL4*, *IL10*, *IFNG* and *TNF* were quantified in biopsies of skin lesions from human patients with localized (LCL) or diffuse cutaneous leishmaniasis (DCL) or in biopsies of normal skin from healthy controls. Statistical analysis was performed using One-way ANOVA followed by Tukey’s multiple comparisons test. Level of significance: p > 0.05 (ns, not significant); *p < 0.05; **p < 0.01; ***p < 0.001; ****p < 0.0001.

## DISCUSSION

Two major challenges in CL are (a) the still incomplete knowledge of the mechanisms leading to non-healing CL and (b) the limited number of therapeutic options. With respect to the pathogenesis of chronic CL, the dual role of macrophages as host and effector cells as well as the bipartite metabolism of L-arginine via the ARG1 and the NOS2 pathways has received particular attention, but the heterogeneity of macrophages and the spatiotemporal regulation of ARG1 and NOS2 are a matter of ongoing intense investigation (*5*). As to the treatment of chronic CL, only few antiparasitic drugs for local or systemic application are currently in clinical use, all of which have significant side effects and limited efficacy and are often not available in endemic countries (*46–48*). Despite the discovery of many new compounds with promising antileishmanial activity (*49*), none of them has been approved for clinical application to date. The present study offers new insights into both areas of CL research. *In vivo* infection experiments with transgenic mouse lines and *ex vivo* immunological, metabolomic and scRNAseq analyses clearly demonstrated that ARG1 accounts for the chronic course of CL in *L. mexicana*-infected C57BL/6 mice. Unlike WT controls and *Arg2*-deficient mice, mice lacking *Arg1* in all hematopoietic or endothelial cells (Arg1ΔTek) or only in monocytes/macrophages (Arg1ΔCx3cr1) were able to control the parasite and to heal the infection. Mechanistically, ARG1 expressed by CX3CR1^+^ monocytes, together with IFNγ, promoted the differentiation of monocytes into *Arg1^+^Nos2^+^Cxcl9/10^+^MhcII^+^*inflammatory macrophages (iMACs) that formed the predominant host cell niche for the parasite and caused a long-lasting depletion of L-arginine in the skin lesions, which impeded protective immunity. Finally, we found that oral supplementation of L-arginine not only prevented the development of CL, but was also sufficient to induce clinical resolution of established chronic CL lesions and to reinforce protective Th1 immunity. Importantly, mice that were cured by L-arginine treatment, had a markedly reduced tissue parasite load, did not develop clinical relapses and were resistant to reinfections. We have obtained no evidence that oral L-arginine would feed the arginase of *L. mexicana* or ARG1 of host cells, which is expressed at high levels in chronic mouse and human *L. mexicana* skin lesions. Thus, we suggest oral L-arginine as host-directed therapy also for human *L. mexicana* CL patients.

Previous studies on L-arginine metabolism and the role of arginase in leishmaniasis have mainly focused on the role of ARG1 in *L. major*-infected mice. In *L. major*-infected C57BL/6 mice, which only weakly express ARG1 and show a self-healing course of CL due to an efficient Th1 response, the deletion of *Arg1* did not further accelerate the control of infection (*50*). In contrast, in Th2-prone BALB/c mice, which develop non-healing local and lethal visceral disease after *L. major* infection, disease progression was prevented by pharmaceutical inhibition (*5, 51, 52*) or genetic deletion of *Arg1* in hematopoietic and endothelial cells (*32*). *Arg1* deletion led to a prominent upregulation of NOS2-dependent NO production in the absence of an altered T cell response, but further mechanistic analyses were not performed (*32*). In our present study, we used a fundamentally different infection model (*L. mexicana*-infected C57BL/6 mice) that more closely resembles chronic CL in humans, which is typically characterized by a mixed T (helper) cell and cytokine response after infection with *L. mexicana* or other *Leishmania* species (*34, 53*). Using this model, we found that the expression of ARG1 not only impeded the activity of NOS2, but, together with IFNγ, also promoted the differentiation of newly recruited monocytes into iMACs, which serve as host cells for *Leishmania* (due to their simultaneous expression of ARG1 and NOS2) and accelerate inflammation in the skin (due the release of chemokines). Although IFN-γ induced NOS2 in *L. mexicana*-infected mice, it unexpectedly also appeared to contribute to the development of pathology. Paradoxical activities of IFNγ are not unprecedented. For example, in visceral leishmaniasis of hamsters, IFNγ induced several counterprotective genes including *Arg1* (*54*). In *L. amazonensis* infection of mice, IFNγ mediated the early recruitment of CCR2^+^ monocytes that functioned as parasite permissive host cells (*55*).

The drugs that are currently used for the treatment of chronic CL (pentavalent antimony preparations, miltefosine, amphotericin B, azoles, paromomycin) show highly variable effectiveness, exert a significant degree of toxicity, and, partly, are also expensive. Except for locally applied compounds that promote wound-healing, and measures of simple wound care, which prevents superinfections (*56–58*), host-directed therapies have not yet entered standard clinical use in CL. L-arginine, a semi-essential amino acid, is normally synthesized endogenously by the host, but often becomes limiting during states of high metabolic demand (e.g., during infections, malignant diseases or chronic inflammations) (*27, 28, 59–61*). Therefore, L-arginine has been considered as adjuvant therapeutic agent in patients with severe or chronic infectious diseases. It can be conveniently applied via the oral route and has not shown toxicity even at high doses (*62*). However, previous randomized control trials on the oral application of L-arginine in patients with active tuberculosis yielded heterogenous results with little or no clinical improvement in the L-arginine-treated group (*63–66*). In the *L. major* BALB/c mouse model discussed above, the intraperitoneal injection of low doses of L-arginine (3 x 10 mg/week, starting on day 14 p.i.) only delayed, but did not prevent disease progression during the very short 3-weeks follow-up period (*67*). These rather disappointing results should not discourage further investigations of the use of L-arginine for the treatment of infections with intracellular pathogens as documented by the present study, in which 85% of *L. mexicana*-infected mice with full-blown disease were cured by oral L-arginine (**Fig. 5**, **fig. S7**).

Treatment with L-arginine restored normal levels of L-arginine in infected tissues. This was paralleled by an increased number of CD4^+^ and CD8^+^ T cells in the skin lesions (**Fig. 6F**) and a qualitative shift of the T helper cell response from a mixed Th1/Th2 phenotype with high IL-4 towards a predominance of Th1 cells (**Fig. 6A, B** and **G**). Mechanistically, these effects of L-arginine likely result from the enhanced chemokine expression (**Fig. 6C** and **D**) and the upregulation of IL-12 (**Fig. 6A**), respectively, although direct or intrinsic effects of L-arginine on the proliferation and functional differentiation of T cells observed in other settings (*27, 68–73*) cannot be excluded. Our observations on the redirection of the T cell response by L-arginine are of immediate relevance for its potential application in human patients with CL. Chronic skin lesions of LCL or DCL patients infected with *L. mexicana* or other *Leishmania* species are positive for IFNγ and NOS2 as well as IL-4 and ARG1 (**Fig. 7**) (*34, 53, 74–77*). High levels of IL-4 and ARG1 and a lack of IFNγ are particularly characteristic for DCL lesions and the occurrence of relapses (*45, 75, 77, 78*). In contrast, a strong IFNγ response is an indicator and prerequisite for the healing of CL (*11, 79*), although it also contributes to tissue damage in acute CL (*80*). Considering the limited therapeutic options for chronic LCL or DCL, we propose oral L-arginine as novel host-directed therapy for chronic CL. A phase I/IIa clinical trial, which evaluates L-arginine as an adjunct therapy in addition to a standard treatment in patients with chronic *L. mexicana* infections (LCL or DCL), would be a sensible first approach for the clinical application of our findings.

The present study has several limitations. Except for the *ex vivo* analysis of human *L. mexicana* LCL and DCL lesions, the work was performed in a preclinical model and restricted to *L. mexicana* infections. 15% of the *L. mexicana*-infected mice failed to respond to oral L-arginine for currently unknown reasons. Further work is also required with respect to the minimal dose and duration of L-arginine therapy and the molecular mechanisms underlying the induced Th1 shift. Finally, prior to application of L-arginine in human LCL or DCL patients, it would be desirable to obtain data on the L-arginine concentration in human skin lesions and to carry out more detailed gene expression profiling of CL biopsies comparing patients with different *Leishmania* infections and stages of disease.

In summary, the presented work identified ARG1 as central metabolic pathway that causes chronicity of *L. mexicana* infections following L-arginine depletion in the infected tissues. ARG1 cooperated with IFN-γ in driving a self-perpetuating cycle of monocyte recruitment and development of inflammatory macrophages as host cells for *Leishmania* parasites. Finally, we defined oral L-arginine as a novel host-directed therapeutic strategy that restored arginine availability in the skin, enhanced Th1 immunity, and thereby caused resolution of chronic CL.

## MATERIALS AND METHODS

### Mouse Strains

Female, 6-weeks old C57BL/6J and C57BL/6N mice were purchased from Charles River (Sulzfeld, Germany). Transgenic *Arg*2^-/-^ (Stock No. 020286), *Arg1*^fl/fl^ (Stock No. 008817) and *Tek*cre mice (Stock No. 008863) on a C57BL/6 background were bought from the Jackson Laboratory (Bar Harbor, U.S.A.). *Cx3cr1*cre C57BL/6 mice (stock No. 036395-UCD) were from the MMRRC repository (U.S.A.). *Il10ΔCd4* mice, also on C57BL/6 background, were provided by Prof. Axel Roers (University Hospital Heidelberg, Germany). Conditional *Arg1*-deficient mice (B6 *Arg1ΔTek* and B6 *Arg1ΔCx3cr1*) were generated through eight generations of backcrossing and subsequent inter-crossing thereafter (*32, 36, 44*). Mice used for experiments were between 6 and 14 weeks old. For infection experiments, female KO or transgenic mice and their age-matched WT littermate controls were used. All mice were housed in specific pathogen-free (SPF) conditions in cages with filter tops at the Franz-Penzoldt Preclinical Experimental Animal Center or at the animal facility of the Institute for Clinical Microbiology, Immunology and Hygiene, University Hospital Erlangen. All animal experiments were approved by the regional animal welfare committee of the governments of Middle or Lower Franconia, Germany.

### Parasites and mouse infections

Promastigotes of the *L. mexicana* strain (MNYC/BZ/62/M379) (*81*) were stored in liquid nitrogen in small aliquots. For experiments, one aliquot was thawed and cultured in complete modified Schneider’s medium (prepared and supplemented as described (*16*)) at 28°C with 5% CO_2_ for up to 5 to 7 passages. Stationary-phase promastigotes were harvested by centrifugation at 3,400 x g for 10 min and the number of viable parasites was determined using trypan blue staining and counting in a Neubauer chamber (Brand, Wertheim, Germany). For infection, 3×10^6^ *L. mexicana* promastigotes in 50 μL in PBS (Merck-Millipore, Burlington, MA, USA) were injected subcutaneously into the skin of both hindfeet using a Sub-Q needle (0.30 mm (30G) x 8mm). Clinical disease progression was monitored by weekly measurements of the skin swelling with a metric caliper (Kroeplin, Schlüchtern, Germany) and determining the % increase compared to the thickness prior to infection. The parasite burdens in the skin lesions, dLNs and spleens were determined by limiting dilution (LD) analysis with serial 3-or 5-fold dilutions and 12 replicates per dilution step as described before (*16, 32*).

### Collection of mouse plasma

Blood was obtained from anesthetized mice by cardiac puncture using a 1 mL syringe and collected in 1.5 mL tubes coated with 0.5 M EDTA solution (Sarstedt, Germany). Blood samples were centrifuged at 4°C and 1,500 × g for 10 min. The upper plasma layer was transferred into fresh 1.5 mL Eppendorf tubes and stored at -20°C until further use.

### L-arginine and anti-IFN-γ treatment of mice

A solution of 40g/L L-arginine monohydrochloride (HPLC grade, Sigma-Aldrich) in tap water was prepared, adjusted to a pH of 3.0 and passed through a 0.02 µm sterile filter. For **prophylactic** treatment, the L-arginine-containing drinking water was provided to the mice from day 10 p.i. onwards until the end of the experiment. In the **therapeutic** setting, L-arginine treatment via the drinking water was initiated when the increase of skin swelling had reached ≍100% (full blown disease state). The treatment was either given until the end of the experiment or stopped after clinical healing of the lesions. In both settings, mice were supplied with freshly prepared L-arginine-containing drinking water every 3-4 days to avoid microbial contamination.

For anti-IFN-γ treatment, *L. mexicana*-infected C57BL/6N mice received twice weekly an intraperitoneal injection of 50 µg rat IgG1-anti-mouse IFN-γ monoclonal antibody (mab) in 100 µL PBS (clone XMG1.2, BioXCell). Control mice were treated with an isotype control mab (HRPN Rat IgG1, BioXCell).

### Quantitative real-time PCR (qPCR)

Total RNA was extracted from homogenized mouse tissue using the TRIfast reagent (Peqlab). Subsequently, cDNA was synthesized from 3 μg of RNA utilizing the high-capacity cDNA reverse transcription kit following the manufacturer’s protocol (Thermo Fisher Scientific). Gene expression levels were quantified using a 384-well plate Viia7 Real-Time PCR system and TaqMan® gene expression assays and master mix (Applied Biosystems; Thermo Fisher Scientific). For normalization of the target gene, mouse Hprt-1 (hypoxanthine-guanine phosphoribosyl transferase-1) was used as an endogenous control gene according to the formula 2^-(Ct^ ^target−Ct^ ^endogenous^ ^control)^ × 10^4^ (arbitrary factor). In the case of human samples, Gapdh (glyceraldehyde-3-phosphate dehydrogenase) was utilized. For the gene-specific assays used, see **table S1**.

### Western blots

Naive or infected tissue samples (skin lesions or dLNs) were prepared using 300 µL of ice-cold Triton-X lysis buffer (20 mM Tris-HCl pH 7.8, 150 mM NaCl, 1% Triton-X, 1 mM EDTA, 1 mM PMSF, 1 × protease inhibitor mix [cOmplete EDTA free tablets, Roche]). For skin lesions and dLN samples, a tissue homogenizer (VWR Star-Beater, stainless steel beads, 3 mm diameter) was used for 3 min at a shaking frequency of 30/s. Skin lesions were homogenized twice under the same conditions and then centrifuged at 600 × g for 1 min at 4°C (Eppendorf, centrifuge 5417R). Steel beads were removed, and the samples were centrifuged at 20,817 × g for 10 min at 4°C. Supernatants were transferred into a fresh 1.5 mL Eppendorf tube and sonicated (level 2, 10 sec). Protein concentrations were determined using the DC™ Protein Assay (Biorad). 60-80 μg of protein were diluted in sample buffer, separated by SDS-PAGE, and then transferred to a PVDF membrane (Millipore Immobilon-P, 0.45 μm pore size) using the tank blot technique (1h, 1 A). After transfer, non-specific binding sites were blocked using 5% (w/v) skim milk in TBST buffer (0.1% v/v Tween 20 in Tris-buffered saline) for at least 1h at RT. The membrane was incubated with the primary antibody diluted in 5% (w/v) skim milk in TBST buffer for 1h at RT or overnight at 4°C. After two washing steps for 10 min with TBST, the membrane was exposed to the appropriate peroxidase-coupled secondary antibody for 1h at RT. After three additional washes, antibody-protein complexes were detected using chemiluminescence (ECL Plus Pierce® Western Blotting Substrate, Pierce/Thermo Fisher Scientific) and a digital documentation system (ChemiLux Imager, iNTAS). If needed, the membrane was treated with Restore® Western blot stripping buffer (Thermo Fisher Scientific) according to the manufacturer’s protocol and the blocking and antibody incubation process was repeated. Finally, images were processed using Photoshop CS5 (Adobe Systems) and band intensity was quantified with ImageJ. All antibodies used are listed in **table S2**.

### ELISA

To detect IFNγ, IL-4, and IL-10 in tissue lysate, plasma, and cell culture supernatants, a sandwich ELISA was performed. All washing steps were carried out using wash buffer (0.05% Tween-20 in PBS). A 96-well Maxisorp™ flat-bottom plate (Thermo Fisher Scientific) was coated with purified rat anti-mouse IFNγ, IL-4, or IL-10 antibody diluted in PBS at the appropriate concentration and incubated overnight at 4 °C in a humidified chamber. After three washes with wash buffer, wells were blocked with 150 µL/well of blocking buffer (10% FCS in PBS) for 1 h at room temperature (RT), followed by a single wash. Samples (30–50 µL/well) including tissue lysates, plasma, and culture supernatants, or two-fold serial dilutions of recombinant mouse IFNγ, IL-4, and IL-10 standards were then added and incubated overnight at 4 °C. After four washes, a biotinylated detection antibody (1 µg/mL in PBS/10% FCS) was added for 1 h at RT. Following four additional washes, bound biotinylated antibodies were detected using Streptavidin-HRP (BioLegend, BD OptEIA Substrate Reagent Set) for 1 h at RT. Tetramethylbenzidine (TMB) was used as the substrate, prepared according to the manufacturer’s instructions. After six washes, 50 µL of TMB substrate solution was added per well and incubated in the dark until a deep blue colour appeared in the highest standard concentration. The reaction was stopped by adding 25 µL/well of 2N H₂SO₄, changing the colour from blue to yellow. Optical density (OD) was measured at 450 nm using a spectrophotometer. Cytokine concentrations were determined from standard curves generated for each assay using SoftMax Pro software. All antibodies used are listed in **table S2**.

### LEGENDplex^TM^ assay

The same tissue lysates that were used for Western blot analysis were also subjected to LEGENDplex™ (BioLegend) analysis. Lysates were centrifuged at 20,817 × g for 10 min using an Eppendorf 5417R centrifuge, and the resulting supernatants were collected for analysis. All reagents were prepared according to the manufacturer’s instructions, and the assay procedures were followed as described for V-bottom plates in the LEGENDplex™ Multi-Analyte Flow Assay Kit manual (Cat. No.: 741294 and 741054), except that assay standards, samples and kit reagents were used at half of the volumes specified in the manual. Chemokine levels were quantified using the LEGENDplex™ Mouse Proinflammatory Chemokine Panel 1 (13-plex) following the Flex Protocol V02 (Cat. No. 741294, Lot B429986). Cytokine levels were assessed using the LEGENDplex™ Mouse Th1/Th2 Panel (8-plex) with VbP V03 (Cat. No. 741054, Lot B449447)

### Flow cytometry

Single cell suspensions from skin lesions and dLNs were prepared by cutting the tissues into small pieces and digestion in 2 mL HBSS (containing CaCl_2_ and MgCl_2_; Gibco, Thermo Fisher), 1% FCS, 0.1 mg/mL DNAse I (Roche) and 0.2 mg/mL collagenase P (Roche) at 37°C using a thermo-mixer at 1,400 rpm. The skin cell suspensions were sequentially passed through a 100 μm and 40 μm cell strainer (Corning, Amsterdam, NL) and centrifuged at 1,400 rpm for 10 min. Enrichment of viable skin cells was performed using density gradient centrifugation (Percoll, GE Healthcare). dLN cells were passed through a 40 μm cell strainer (Corning, Amsterdam, NL), washed and resuspended in PBS. Single cell suspensions were stained first with Zombie Aqua Fixable Viability Dye 516 (BioLegend 423102), washed and resuspended in FACS buffer (PBS/ 1% (v/v) FCS/ 2 mM EDTA) and stained with rat-anti-mouse CD16/CD32 for 5 min to block Fc receptors. Fluorophore-coupled surface antibodies against surface antigens (CD3, CD4, CD8, CD11b, Ly6G, NK1.1, SiglecF, TCRγδ) were added at varying concentrations based on prior titration (**table S3**). The cells were incubated in the dark at 4°C for 30 min, and unbound antibodies were washed off. In some cases, biotinylated antibodies were used, and selected fluorophore-coupled streptavidin was incubated for 20 min at 4°C. Measurements were done with the help of a BD LSR Fortessa™ flow cytometer equipped with an ultra-violet (355 nm), blue (488 nm), yellow-green (561 nm), red (640 nm), and violet (605 nm) laser. Cytometer performance was monitored regularly, and compensation controls were carried along in each experiment. FlowJo v.10.8.1 software was used for analysis.

### Restimulation of cells and intracellular antigen staining

For restimulation of dLN cells, 250,000 cells were resuspended in 250 µL RPMI1640 cell culture medium (ThermoFisher) supplemented with 10 % FCS and exposed to medium alone, plate-bound monoclonal anti-CD3/anti-CD28 antibodies or freeze-thaw lysates of *L. mexicana* promastigotes (FT-Lmex) for 72 h. Subsequently, culture supernatants were harvested and analyzed for their cytokine content (e.g., IFN-γ, IL-4, IL-10).

For intracellular cytokine staining, 1-2 × 10^6^ (skin lesion) cells were incubated in a round bottom tube in 200 µL RPMI1640 cell culture medium with 5% FCS and supplemented with 2µM monensin (Biolegend, cat: 420701, lot: B333269) for 4 hours at 37°C. Cells were then washed and stained for surface antigens (**table S3**). Cells were fixed using the ebioscience^TM^ FoxP3/transcription factor staining buffer set (Thermo Fisher Scientific) or CytoPerm fixation buffer (BD Biosciences), depending on the intracellular antigen. CytoPerm/CytoFix fixation was used for intracellular staining of IFN-γ, while FoxP3/Transcription Factor staining buffer set was used for intracellular staining of ARG1 and NOS2. Appropriate concentrations of fluorophore-coupled antibodies (ARG1, NOS2, and IFNγ) were added and incubated at 4°C in the dark overnight. The next day, unbound antibodies were washed off and analyzed with a BD LSRFortessa™ flow cytometer.

### Sorting of ARG1^+^NOS2^+^ macrophages

Single cell suspensions were stained for surface antigens (CD11b and SiglecF) and intracellular staining (ARG1 and NOS2) was performed (**table S3**). Afterwards, cells were washed with FACS buffer (PBS/ 1% (v/v) FCS/ 2 mM EDTA) and passed through a 50 μm filter. ARG1^+^NOS2^+^ cells were sorted with the cell sorter MoFlo Astrios EQI. The gating strategy is detailed in **fig. S3D**.

### Immunohistology and confocal laser scanning fluorescence microscopy (CLSFM)

Using a cryotome (CryoStar NX70, Thermo Fisher Scientific) 5 μm thick frozen sections from skin tissue were prepared, embedded in TissueTek® (Sakura Finetek, Germany), and transferred to StarFrost® adhesive slides (Knittel Glasbearbeitungs GmbH, Germany). After a 1 h drying period, sections were fixed with acetone (-20°C, 10 min), dried again for 20 min, and surrounded with liquid blocker (PAP-PEN®, BD Biosciences). After rehydration with PBS/ 0.01% (v/v) Tween 20 for 20 min, non-specific binding sites were blocked with blocking buffer (5% normal donkey serum, 0.1% Saponin in PBS) for 20 minutes. Tissue sections were incubated with primary antibodies (**table S4**) diluted in PBS/ 0.5% (v/v) normal donkey serum for 1 h at RT or, in the case of ARG1 staining, at 4°C overnight. Following three washing steps, the sections were incubated with fluorochrome-conjugated secondary antibodies (**table S4**) diluted in PBS/ 0.5% (v/v) normal donkey serum for 1 h at RT. Again, sections were washed three times for 5 min and nuclei were stained with DAPI (4’,6-Diamidino-2-phenylindole, Sigma-Aldrich) followed by extensive washing steps. For the detection of NO, unfixed frozen tissue sections (5 μm) were mounted on adhesion slides (StarFrost®) and treated with PBS/0.01% (v/v) Tween 20 for 15 min. Then, they were incubated with 10 mM 1,2-diaminoanthraquinone (DAQ, Sigma-Aldrich) in PBS/10% (v/v) DMSO at 37°C for 45 min, followed by three 5-min washing steps and DAPI staining for 30 minutes at RT. After another round of three 5-min washings, the slides were mounted using ProLong™ Diamond Antifade Mountant (Invitrogen) and allowed to dry for at least 12 h at 4°C in the dark before analysis within 1-2 days using a Zeiss LSM700 microscope (4 laser system, 405 nm, 488 nm, 555 nm, 639 nm). Image acquisition, processing and analyses were performed using ZEN 3.0 (blue edition). To ensure accuracy, proper controls were used to exclude non-specific binding of primary and secondary antibodies. For directly conjugated antibodies, a corresponding isotype control was included. The specificity of the anti-nitrotyrosine antibody (for the indirect measurement of NO levels in the tissue) was verified using 10 mM 3-nitro-L-tyrosine for blocking of the anti-nitro-tyrosine antibody (Sigma-Aldrich).

### Detection of *L. mexicana* in sorted ARG1^+^NOS2^+^ cells by immunofluorescence microscopy

Sorted ARG1^+^NOS2^+^ cells were centrifuged at 1,400 rpm for 10 min, the supernatant was removed and the cell pellet was resuspended in 200 µL of PBS The suspension was split into two halves: one for *L. mexicana* staining and the other for the secondary-antibody-only control. For staining, 50-100 µL of cell suspension was transferred to an adhesion slide and allowed to adhere for 1 h. The human anti-*L. mexicana* serum, diluted in FACS saponin buffer (0.5% saponin, 2% FCS in double distilled water), was added and incubated for 2 h at RT. The cells were washed twice with FACS saponin buffer and incubated with anti-human FITC-conjugated secondary antibody diluted in FACS saponin buffer for 1-2 h at RT. The cells were washed twice for 5 min and mounted with DAPI Fluoromount-G® (Southern Biotech). Slides were kept overnight at RT and images were taken using a KEYENCE fluorescence microscope (BZ-X800/ BZ-X810) with a 40× objective.

### Single-cell RNA-sequencing (scRNA-seq)

Skin lesions from *Arg1*^fl/fl^ and *Arg1ΔCx3cr1* mice were digested (2 mL HBSS containing CaCl_2_ and MgCl_2_ [Gibco, Sigma-Aldrich], 1% FCS, 0.1 mg/mL DNAse I [Roche] and 0.2 mg/mL collagenase P [Roche]; 37°C thermomixer, 1,400 rpm), sequentially passed through 100 µm and 40 µm cell strainers (Corning, Amsterdam, NL) and centrifuged at 1,400 rpm for 10 min. Enrichment of viable cells was performed using density gradient centrifugation (Percoll, GE Healthcare). Dead cells and free *Leishmania* were removed using MACS Dead Cell Removal Kit (Miltenyi Biotec) according to the manufacturer’s instruction. Next, cDNA synthesis, barcoding and library preparation were performed using Chromium Next GEM Single Cell 3’ Reagent Kit v.3.1 from 10x Genomics. The libraries were sequenced on an Illumina HiSeq 2500 (Illumina, San Diego) with a read length of 100 bp for read 1 (cell barcode and unique molecule identifier [UMI]), an 8 bp i7 index read (sample barcode), and 100 bp for read 2 (RNA sequence). The reads were demultiplexed using Cell Ranger (v4.0, 10x Genomics) ‘mkfastq’ in combination with ‘bcl2fastq’ (Illumina). Sequencing of the *Arg1^f^*^l/fl^ and *Arg1ΔCx3cr1* flow cells yielded approximately 255 million reads.

### ScRNA-seq data analysis

A customized reference database was constructed using the *L. mexicana* genome (Genome assembly ASM23466v4, Wellcome Trust Sanger Institute) in conjunction with the *Mus musculus* genome (GRCm38, modified to align with Cell Ranger requirements). Single cell RNA sequencing reads derived from WT (*Arg1*^fl/fl^) and KO (*Arg1ΔCx3cr1*) skin lesion cells were quantified against this reference database using Cell Ranger (v4.0, 10x Genomics). This analysis identified in total 3,845 cells (WT: 2,101, KO: 1,744), along with 40,534 gene features. Data preprocessing and analysis was conducted using the Seurat package v4. Cells with a mitochondrial gene content greater than 10% or gene features with fewer than 4000 counts were excluded from subsequent analyses. A global normalization method was applied through the “LogNormalization” function, followed by data scaling within Seurat. The “vst” method identified 2,000 variable features, which were used for principal component analysis (PCA) and further downstream analyses. Cell clustering was performed using the Louvain algorithm at a resolution of 0.8. Clusters were visualized using the Uniform Manifold Approximation and Projection (UMAP) method. Pseudotime analysis to create cell differentiation trajectories was conducted with Monocle 3. Additionally, RNA velocity dynamics were explored using the scVelo tool. To detect cells infected with *Leishmania*, a specific gene expression module was defined for *Leishmania* genes. Expression scores were visualized using a feature plot, and the “AddModuleScore” function within Seurat was employed to identify clusters positive for *Leishmania* infection.

### LC-MS/MS metabolites analysis of tissue samples

Tissue samples (skin lesions or dLNs, at least 30 mg) were thoroughly homogenized using an appropriate homogenization system (Precellys Kit, Bertin Technology, France). Ethanol/phosphate buffer (85:15 v/v, 3 μL buffer per 1 mg tissue) was used as extraction buffer. The samples were then centrifuged at 5,000 × g for 5 minutes at 4°C. To determine amino acids and biogenic amines in tissue supernatant, the respective samples were repeatedly dried under nitrogen air flow and reconstituted with a derivatization buffer consisting of equal parts pyridine, ethanol and water, as previously described (*82*). Derivatization was performed with 5% phenyl isothiocyanate (PITC) for 30 min at RT. The samples were then dried again and dissolved in a methanol solution containing 5 mM ammonium acetate. Separation and quantitation of the resulting phenylthiocarbamyl derivatives were carried out using reverse-phase liquid chromatography coupled with an ESI-MS/MS system in positive ionization mode (API6500+, Sciex Darmstadt, Germany; HPLC 1260 series, Agilent Technologies, Waldbronn, Germany). The liquid chromatography utilized an Agilent Eclipse XDB-C18 column (3.5 µm, 3×100 mm) as the stationary phase. The mobile phase consisted of two solutions: 0.2% formic acid in water and 0.2% formic acid in acetonitrile, using a gradient ramping towards higher acetonitrile content. The transitions, along with the matching deuterated internal standards for quantitation in MRM mode, are detailed in **table S5**. Statistical, and pathway analyses were performed using MetaboAnalyst 5.0.

### LC-MS/MS metabolites analysis in plasma

Samples, calibrators and quality controls (25 µL each) were pipetted into a deep well plate (Greiner BioOne, Germany), followed by the addition of an internal standard (IS) mix (50 µL). Precipitation reagent (400 µL) was then added, and the samples were mixed for 30 s before centrifugation at 4,000 × g for 5 minutes to precipitate proteins. The supernatant was transferred to a 96-well deep well plate. Subsequently, 5 µL of the supernatant was injected into an API6500 mass spectrometer using an Agilent system with an autosampler (API6500+, Sciex Darmstadt, Germany, HPLC1260 series, Agilent Technologies, Waldbronn, Germany). Gradient chromatography was performed on the analytical column provided with the kit, maintained at 35°C. The initial conditions were set to 100% mobile phase A, with a flow rate of 800 mL/min, as specified in the manufacturer’s package leaflet. Mass spectrometer settings included an ion spray voltage of 3500 V, a source temperature of 400°C, a curtain gas setting of 45, low collision gas, source gas 1 at 35, and source gas 2 at 45. Collision energy was optimized for each analyte (**table S5**). Data acquisition was performed in MRM mode using positive ionization for all analytes, and processed with Multiquant software (Sciex Darmstadt, Germany). A four-point calibration curve was used for all analytes, with a total analysis time of 20 minutes. Additional method details, including calibrator concentrations, were provided in the manufacturer’s package leaflet.

### Statistical analysis

Statistical analyses were conducted using GraphPad Prism version 9. Prior to performing statistical tests, datasets were tested for Gaussian distribution, and outliers were identified using the ROUT method. For comparisons between two groups, a two-tailed nonparametric Mann-Whitney U test (if n ≥ 4 samples) or parametric unpaired t test (if n ≤ 3) were applied. For data following a normal distribution from two or more groups, a One-way ANOVA was conducted, followed by Tukey’s multiple comparisons test. Additionally, comparisons between the two groups of the time-course experiments were performed using a mixed-model two-way repeated-measures ANOVA. A p-value of ≤ 0.05 was considered statistically significant. Additionally, Pearson correlation analysis was performed using linear regression. The line fitting was established with a 95% confidence band, and the goodness of fit (R²) was calculated along with the corresponding p-value.

## Supporting information

\\nas-fa0efsusr1\raibu\Desktop\Text

## Abbreviations

*Arg1*/ARG1: arginase 1
CLSFM: confocal laser scanning fluorescence microscopy
DC: dendritic cell
dLN: draining (popliteal) lymph node
dpi: days post infection
IFN: interferon
IL: interleukin
KO: knockout
LD: limiting dilution
NK: natural killer cell
NO: nitric oxide
*Nos2*/NOS2: type 2 (or inducible) nitric oxide synthase p.i., post infection
scRNA-seq: single-cell RNA sequencing
Th: T helper lymphocytes
WT: wild-type.

## Data availability

scRNA-seq data generated during this study have been deposited at the Gene Expression Omnibus under the accession number GSE306503. All other datasets (flow cytometry dataset and metabolomics dataset) are available upon request. Requests should be sent to the corresponding authors, Ulrike Schleicher and Christian Bogdan. For ScRNA-seq analysis, reads were mapped to the mouse genome (refdata-gex-mm10-2020-A) and *L. mexicana* genome (*Leishmania mexicana* MHOM/GT/2001/U1103). For metabolomics analysis, Metaboanalyst (https://www.metaboanalyst.ca/) and for gene set enrichment analysis (https://www.gsea-msigdb.org/gsea/index.jsp) was utilized.

## Code availability

All custom code utilized for the analysis of these data is available upon request as not all datasets are publicly available. Requests should be sent to the corresponding authors, Ulrike Schleicher and Christian Bogdan, or to Baplu Rai.

## LIST OF SUPPLEMENTARY MATERIALS

### Supplementary figures

**Fig. S1:** Induction of ARG1 and the non-resolving phenotype were dependent on CD4^+^ T cell-derived IL-10.

**Fig. S2:** Clinical severity of disease and parasite burdens were comparable between WT and *Arg1* KO mice on day 40-45 p.i.

**Fig. S3:** WT mice exhibited elevated chemokine expression at the site of infection.

**Fig. S4:** Gene expression kinetics and GSEA analysis of *Cd4⁺Ifng⁺* T cell clusters.

**Fig. S5:** The amino acid microenvironment showed significant changes during the progression to chronic CL.

**Fig. S6:** Restoration of L-arginine levels in the tissue following oral L-arginine supplementation correlated with reduced parasite burden.

**Fig. S7:** L-arginine treatment caused resolution of chronic CL lesions and conferred long-lasting immunity.

**Fig. S8:** Th1 response is slightly enhanced in the dLN of L-arginine-treated mice.

### Supplementary tables

**Table S1:** Gene-specific assays for quantitative real-time PCR

**Table S2:** Antibodies used for Western Blot and ELISA

**Table S3:** Antibodies used for flow cytometry and cell sorting

**Table S4:** Antibodies used for CLSFM

**Table S5:** Metabolomic analyses by LC/MS

## AUTHORS CONTRIBUTIONS

Conceived the study and designed the experiments: B.R., U.S., C.B.

Performed the experiments: B.R., A.D., H.S., S.K.

Analyzed the data: B.R., D.B., C.L., H.S., A.D., S.K., M.R., C.B., U.S.

Provided reagents or samples: S.R., I.B.

Provided methodological or technical advice: M.G., C.L., D.B., T.G., M.K., A.B. J.M.

Acquired funding and supervised the project: U.S. and C.B.

Wrote the draft and final version of the manuscript: B.R., U.S. and C.B.

All authors: reviewed, edited, and approved the final manuscript.

## ACKNOWLEDGEMENTS

We are grateful to Dr. Arif Ekici as well as to Uwe Appelt and Markus Mroz from the University Hospital Erlangen core units “Next Generation Sequencing” and “Cell Sorting and Immunomonitoring”, respectively, for their technical support, and to personnel of the Franz-Penzoldt Preclincal Experimental Animal Center for their professional mouse care. We acknowledge Dr. Philipp Tripal and Dr. Benjamin Schmid (OICE) for assistance with image analysis, and Daniel Köppl (Kinderklinik, Universitätsklinikum Erlangen) for support with LC-MS data curation and analysis. We further thank Dr. Baldomero Sanchez-Barragán (UJAT Tabasco, Mexico) for facilitating access to patient samples.

This study was conducted in partial fulfillment of the doctoral theses of Baplu Rai (Dr. rer. nat.) and Sinan Kirici (Dr. med.).

## FUNDING

This research was supported by the German Research Foundation (DFG; RTG 2740 “ImmunoMicroTope”, project number 447268119, projects A1 to A.B., A6 to U.S., and B2 to C.B. and M.K.; CRC1181, project C04 to U.S., J.M. and C.B.; SPP1937 “Innate lymphoid cells”, grants SCHL 615/1-1 and 1-2 to U.S. and grants BO 996/5-1 and 5-2 to C.B.; grants BO996/7-1 and SCHL615/3-1 to C.B. and U.S), the Interdisciplinary Center for Clinical Research (IZKF) of the Universitätsklinikum Erlangen (project A87 to U.S.), the STAEDTLER Foundation Nürnberg (grant to C.B. and J.M.), and by the Programa de Apoyo a Proyectos de Investigación e Innovación Tecnológica of the Universidad Nacional Autónoma de México (grant UNAM-PAPIIT IG2000924 to IB).

## ETHICS STATEMENT

The sampling and analysis of human skin specimens received approval from the Ethics and Research Committees of the Facultad de Medicina, UNAM (Universidad Nacional Autónoma de México) with reference number FM/DI/071/2023. We adhered strictly to the guidelines established by the Ministry of Health in Mexico. All participants and healthy donors (controls) were informed and provided written consent to participate in the study.

## CONFLICT OF INTEREST

The authors declare no competing financial interest in relation to the work described.

## Notes

### Competing Interest Statement

The authors have declared no competing interest.

