## Supplementary material for "Oral L-arginine cures arginase 1-dependent chronic cutaneous leishmaniasis by redirecting the T helper cell response": \\nas-fa0efsusr1\raibu\Desktop\Text

Baplu Rai et al.

Corresponding and senior authors:

Priv. Doz. Dr. Ulrike Schleicher and Prof. Dr. Christian Bogdan, Mikrobiologisches Institut, Universitätsklinikum Erlangen, Wasserturmstraße 3/5, D-91054 Erlangen, Germany;**;**

**This PDF file includes:**

Figs. S1 to S8

Tables S1 to S5

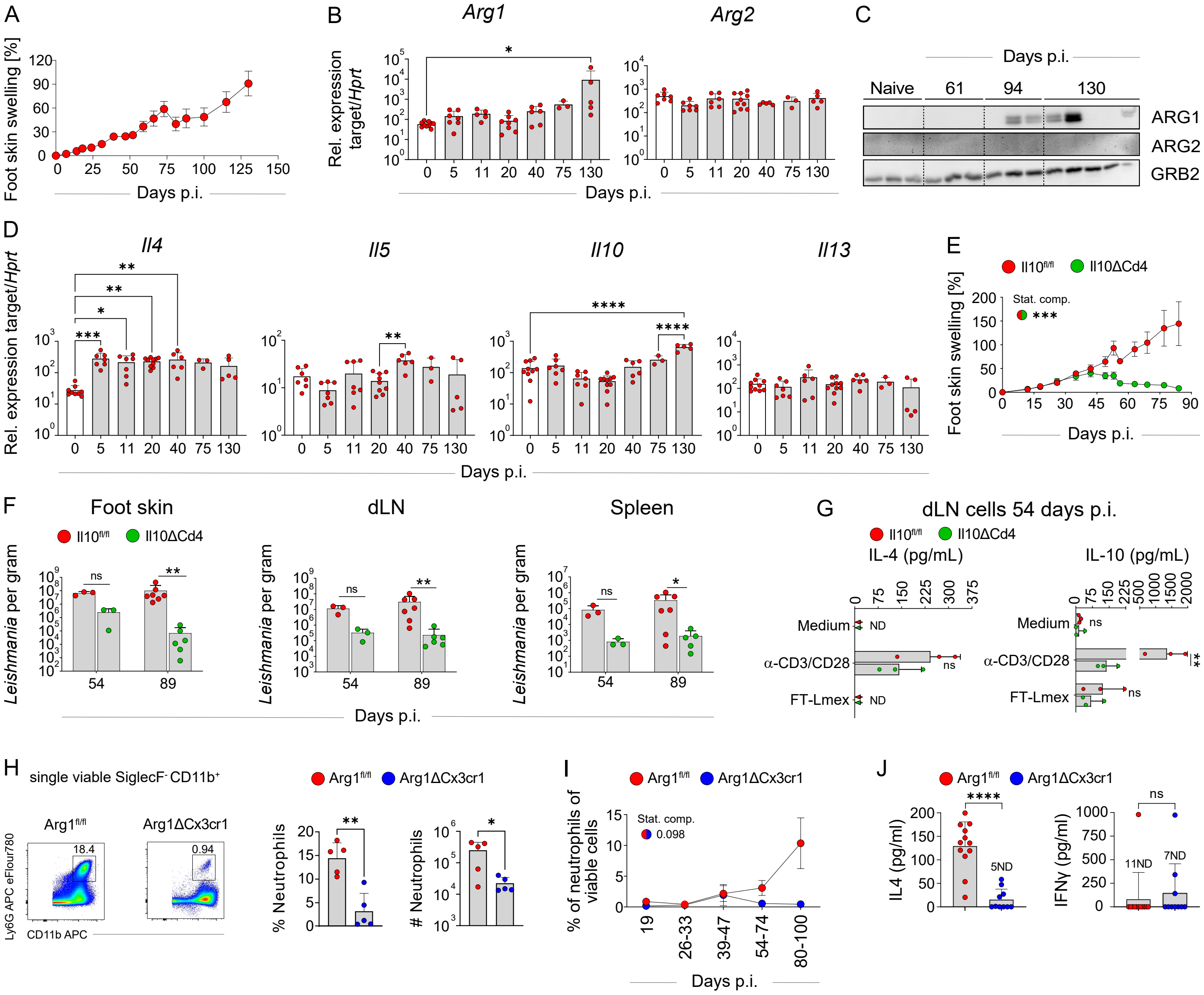


**Fig. S1: Induction of ARG1 and the non-resolving phenotype were dependent on CD4$^+$ T cell-derived IL-10. (A, E)** Skin lesion development in C57BL/6N, *Il10ΔCd4* and *Il10$^{fl/fl}$* mice infected with *L. mexicana.* One of 3-4 independent experiments is shown. **(B, D)** mRNA expression in dLNs at indicated days p.i. as measured by RT-qPCR. Bars show mean ± SD of 3-10 mice per time-point of 3 independent experiments. **(C)** Representative Western blot of ARG1, and ARG2 in dLN lysates (80 µg protein/lane) at indicated days p.i. GRB2: growth factor receptor bound protein-2 as loading control. One of three independent experiments is shown. **(F)** Parasite numbers in the skin lesions, dLNs, and spleens at days 54 and 89 p.i. were quantified by LD analysis in two independent experiments with 3-6 mice per group and time point. **(G)** Supernatants from dLN cells (harvested at days 54 p.i.) were collected after 72 hours of stimulation with medium alone, plate-bound anti-CD3/CD28 or *L. mexicana* freeze-thawed lysate (FT-Lmex). IL-4 and IL-10 contents were analyzed by ELISA. Bars show mean ± SD of 3 mice per group. **(H)** Flow cytometry of skin lesions at day 90 p.i. gated for neutrophils (SiglecF$^-$Ly6G$^+$CD11b$^+$) within single, viable cells. Representative FACS plot from five independent experiments. Skin lesion samples were pooled from 3-4 mice/group. Bar graph showing percentage and absolute numbers of neutrophils that were calculated from 5 independent experiments using FlowJo software. **(I)** Percentage of neutrophils within viable skin cells at different days p.i.. Comparisons between respective groups were performed using a mixed-model two-way repeated measures ANOVA. **(J)** IL4 and IFN-γ level were measured in skin lysates samples at 90 days p.i. using ELISA. Bars show mean ± SD of 10-12 mice per time-point of 2 independent experiments. Statistical significance was determined using two-tailed Mann-Whitney U test **(F, H, J)** or One-Way ANOVA followed by Tukey's multiple comparisons correction **(B, D, G)**. Comparisons between respective mouse groups were performed using a mixed-model two-way repeated measures ANOVA **(E)**. Statistical significance between groups is illustrated using half-circles with the colours of the compared groups **(E)**. Significance levels indicated by asterisks (*) according to the P value. ND, not determined; ns, not significant. $p > 0.05$; *$p < 0.05$; **$p < 0.01$; ***$p < 0.001$; ****$p < 0.0001$.

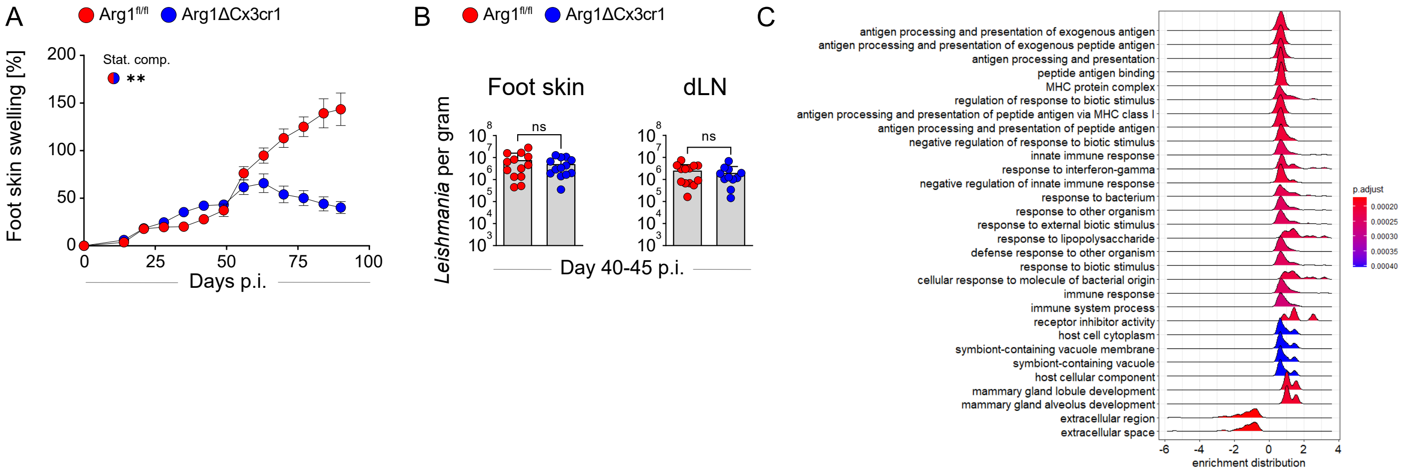


**Fig. S2: Clinical severity of disease and parasite burdens were comparable between WT and *Arg1* KO mice at day 40-45 p.i. (A)** Skin lesion development in *Arg1$^{fl/fl}$* and *Arg1ΔCx3cr1* mice infected with *L. mexicana*. One of 6 independent experiments is shown. **(B)** Parasite numbers in the skin lesion and dLN at day 40-45 p.i. were quantified by LD analysis. Bars show mean ± SD of 14 mice per group from 3 independent experiments. **(C)** Gene Set Enrichment Analysis (GSEA) of WT enriched Arg1$^{+}$Nos2$^{+}$ macrophages clusters (m0-m3, iMAC) from the single-cell RNA sequencing (scRNA-seq) dataset. **(A)** Comparisons between respective mouse groups were performed using a mixed-model two-way repeated measures ANOVA. **(B)** Statistical significance was determined using two-tailed Mann-Whitney U test**.** Significance levels are indicated by asterisks (*) according to the P value. ns (not significant), p > 0.05; *p < 0.05; **p < 0.01; ***p < 0.001; ****p < 0.0001.

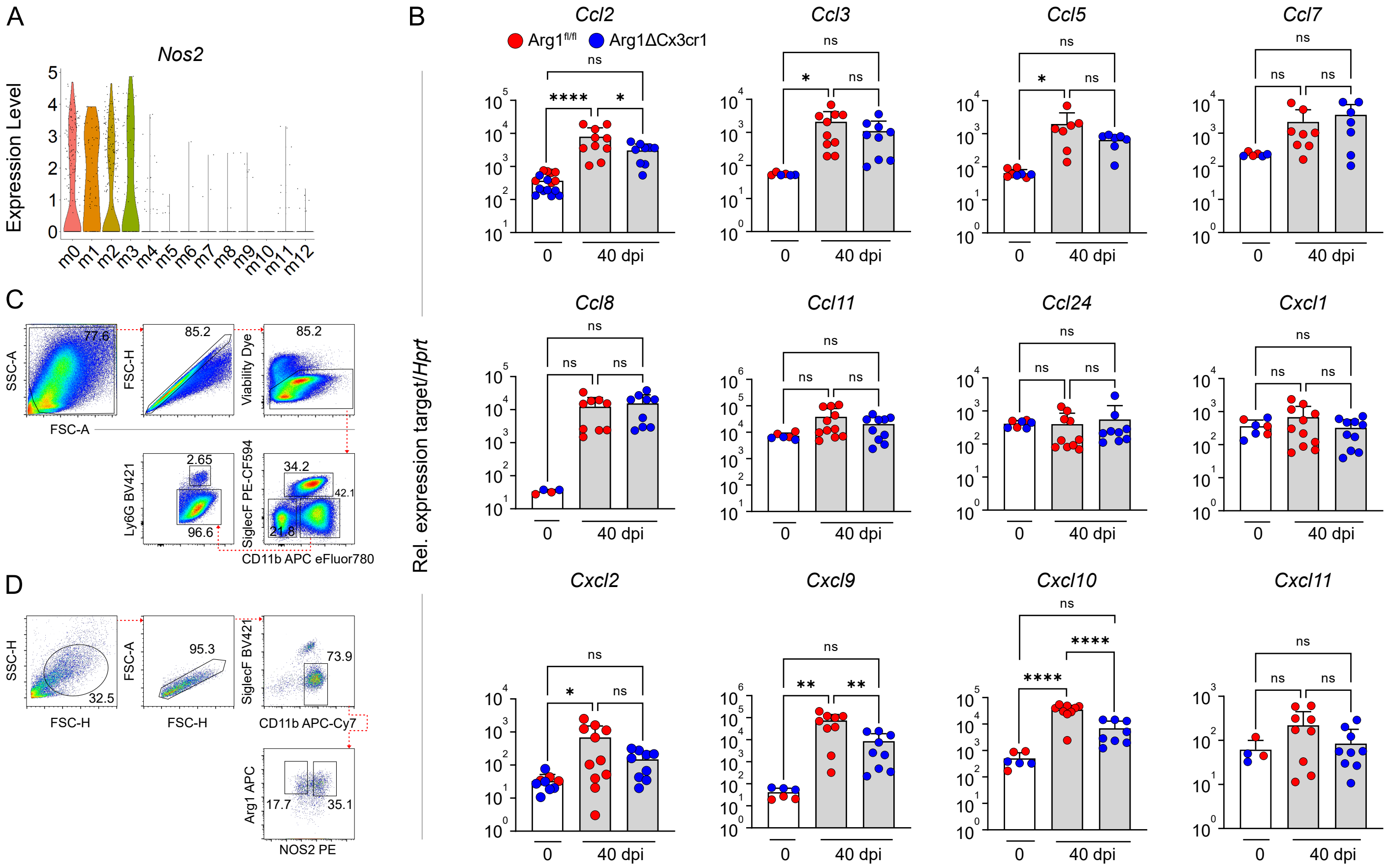


**Fig. S3: WT mice exhibited elevated chemokine expression at the site of infection. (A)** Violin plots depicting *Nos2* expression across myeloid clusters (m0 - m12) from scRNA-seq (10x Genomics) of viable cells pooled from skin lesions (3 mice per group) of *Arg1ΔCx3cr1* (KO) and *Arg1$^{fl/fl}$* (WT) littermates at day 40 dpi. **(B)** Chemokine mRNA expression levels in skin lesions were analyzed by RT-qPCR. Bars represent mean ± SD of 4-15 mice per group from three independent experiments. Statistical significance was determined using One-way ANOVA followed by Tukey's multiple comparisons correction. ns, not significant; significance thresholds: $p > 0.05$ (ns); *$p < 0.05$; **$p < 0.01$; ***$p < 0.001$; ****$p < 0.0001$. **(C)** Myeloid gating strategy used for the analysis of flow cytometry data. **(D)** Gating strategy used to sort ARG1$^+$NOS2$^+$ cells from skin lesions of *L. mexicana*-infected wild-type (WT) mice at 41 days p.i. Sorted ARG1$^+$NOS2$^+$ cells were subsequently analyzed by immunofluorescence staining to visualize *L. mexicana*-infected cells.

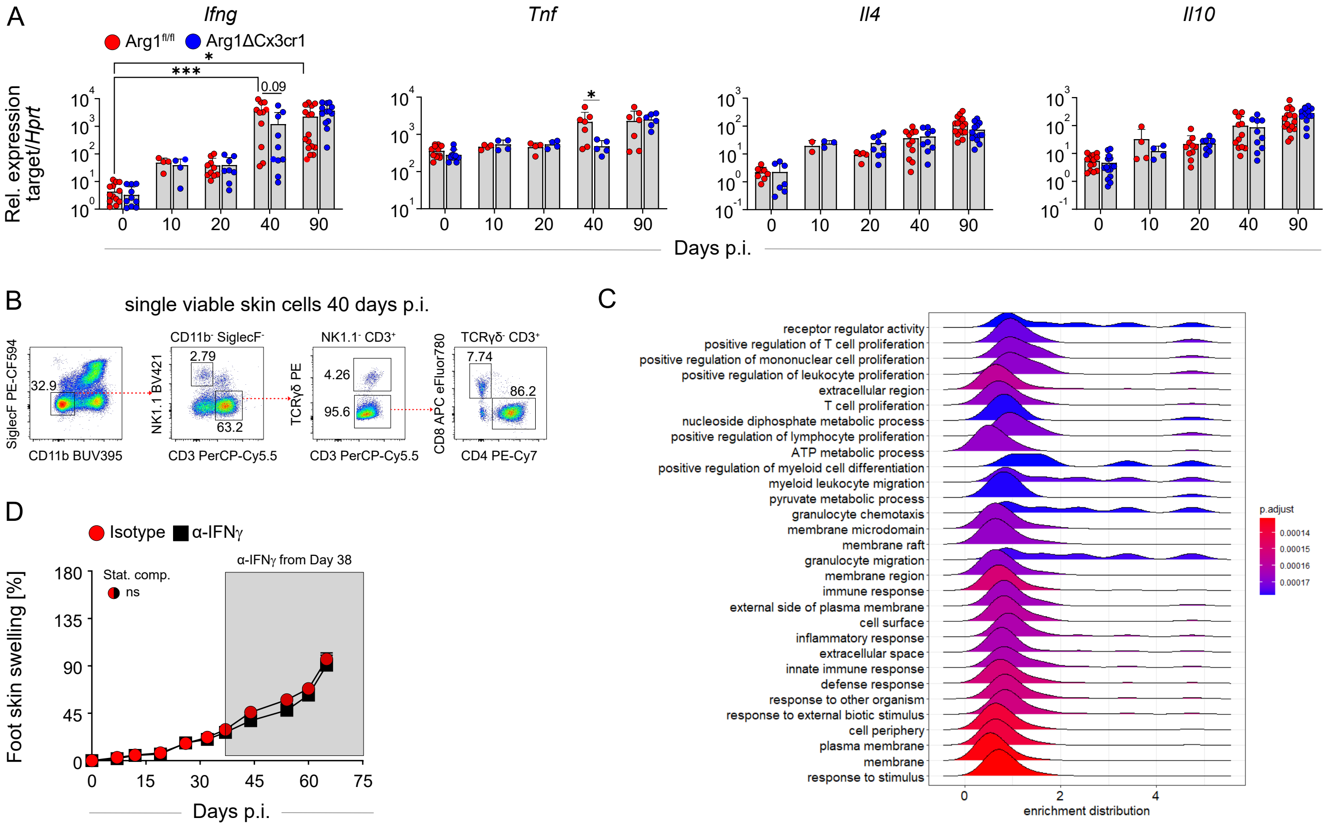


**Fig. S4: Gene expression kinetics and GSEA analysis of *Cd4*⁺*Ifng*⁺ T cell clusters**. **(A)** Time-course analysis of mRNA expression levels of Th1 and Th2 cytokines in skin lesions of Arg1ΔCx3cr1 and WT (Arg1fl/fl) littermate controls at different days p.i., analyzed by RT-qPCR. Bars represent mean ± SD of 4-15 mice per group from four independent experiments. Statistical significance was determined using the One-Way ANOVA followed by Tukey's multiple comparisons correction. **(B)** T cell gating strategy used for analysis of skin lesions at day 40 p.i. **(C)** Gene set enrichment analysis (GSEA) of WT-enriched *Cd4*⁺ *Ifng*⁺ T cell clusters. **(D)** Skin lesion development in anti-IFN-γ- or isotype-treated B6N WT mice. Comparisons between respective mouse groups were performed using a mixed-model two-way repeated measures ANOVA. Statistical significance between groups is illustrated using half-circles with the colours of the compared groups. Significance levels are indicated according to the P value. ns, not significant; significance thresholds: $p > 0.05$ (ns); *$p < 0.05$; **$p < 0.01$; ***$p < 0.001$; ****$p < 0.0001$.

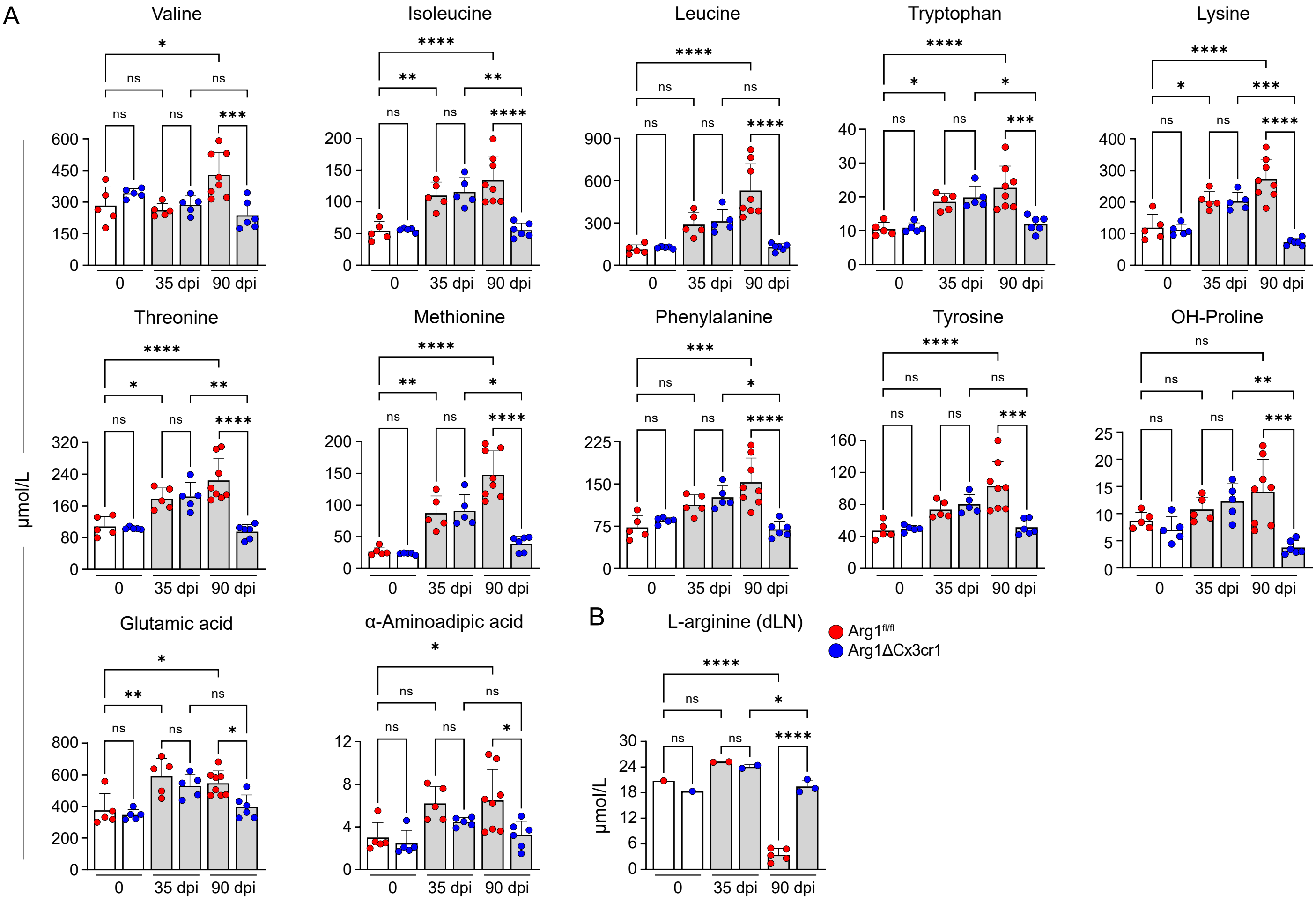


**Fig. S5: The amino acid microenvironment showed significant changes during the progression to chronic CL. (A)** Quantification of metabolite levels at the infection site in *Arg1ΔCx3cr1* and *Arg1$^{fl/fl}$* control mice at different time-points of infection. Data are presented as mean ± SD from 5-8 mice per group. **(B)** Quantification of L-arginine levels in dLNs at different days after infection. LN samples were pooled as follows: day 0, 9-12 mice per datapoint per group; day 35, 2-3 mice per datapoint per group [in total 5 mice per group]; day 90, 1-3 mice per datapoint per group [in total 8 WT/6 KO mice] (n= 1 experiment). Statistical analyses were performed using One-way ANOVA followed by Tukey's multiple comparisons test. ns, not significant; $p > 0.05$; *$p < 0.05$; **$p < 0.01$; ***$p < 0.001$; ****$p < 0.0001$.

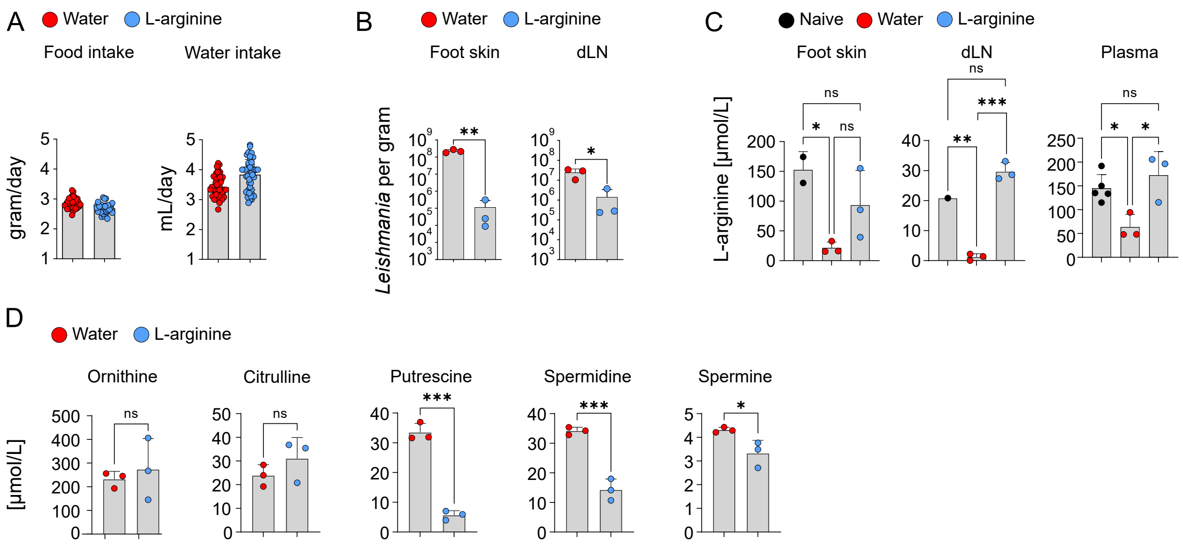


**Fig. S6: Restoration of L-arginine levels in the tissue following oral L-arginine supplementation correlated with reduced parasite burden.** **(A)** Daily food and water intake were monitored in mice on L-arginine-containing drinking water vs. normal water. Bar graphs display average consumption per mouse, reported as grams of food or mL of water intake per day. **(B)** Parasite burden in skin lesions and dLNs were quantified on day 107 p.i. using LD analysis. L-arginine supplementation (40 g/L in drinking water) was started on day 10 p.i. **(C)** L-arginine levels in skin lesions, dLNs and plasma were determined by LC-MS on day 107 p.i. in L-arginine-treated and control mice (on normal water) and compared to levels in naive mice. **(D)** Metabolite levels at the site of infection were quantified using LC-MS in L-arginine- treated and control mice (on normal water) at day 107 p.i. Data are presented as mean ± SD from 3 mice per group. Statistical analysis was performed using parametric unpaired t test **(B, D)** or One-way ANOVA followed by Tukey's multiple comparisons test **(C)**. Significance levels: $p > 0.05$ (ns, not significant); *$p < 0.05$; **$p < 0.01$; ***$p < 0.001$; ****$p < 0.0001$.

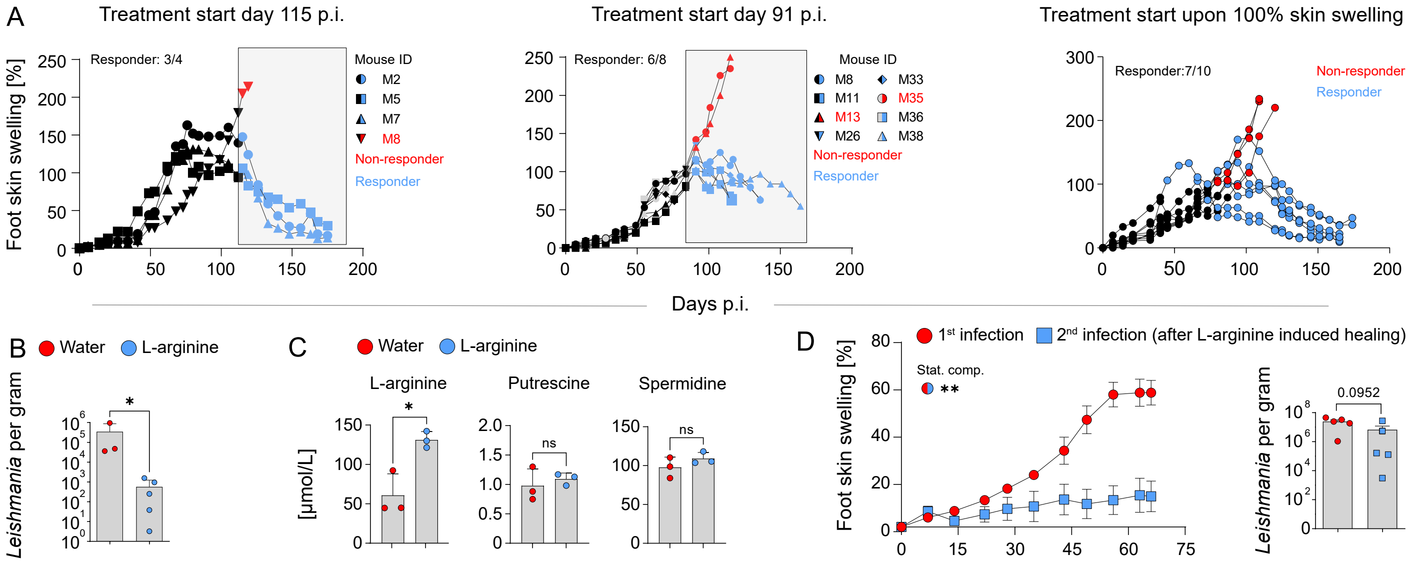


**Fig. S7: L-arginine treatment caused resolution of chronic CL lesions and conferred long-lasting immunity. (A)** Skin lesion development in *L. mexicana*-infected C57BL/6N mice treated with L-arginine (40 g/L in drinking water) compared to untreated (water) controls. L-arginine treatment was initiated, when an increase of the skin swelling of ≥ 100% was reached. Data from individual mice across three independent experiments are shown. Blue lines represent responders; red lines indicate non-responders. **(B)** Parasite burdens in skin lesions of L-arginine-treated mice (75 - 100 days after start of treatment) and controls on normal drinking water (166 days p.i.) were quantified by LD analysis. **(C)** Plasma levels of L-arginine, putrescine and spermidine were measured by LC-MS in mice 10 days after initiation of L-arginine treatment. Three mice per group were analyzed. **(D)** Skin lesion development after reinfection of previously healed mice with *L. mexicana* (three mice were from a therapeutic and two mice from prophylactic regimen). 6-weeks-old naive C57BL/6N mice served as controls and were infected with the same batch of *L. mexicana* promastigotes. Parasite burden in skin lesions was quantified at 67 days p.i. by LD analysis (right panel). Statistical significance was determined using a two-tailed Mann-Whitney U test **(B, D)** or parametric unpaired t test **(C)**. Comparisons between respective mouse groups were performed using a mixed-model two-way repeated measures ANOVA. Statistical significance between groups is illustrated using half-circles with colours of the compared groups **(D)**. Significance levels are indicated by asterisks (*) according to the P value. Levels of significance: p > 0.05 (ns, not significant); *p < 0.05; **p < 0.01; ***p < 0.001; ****p < 0.0001.

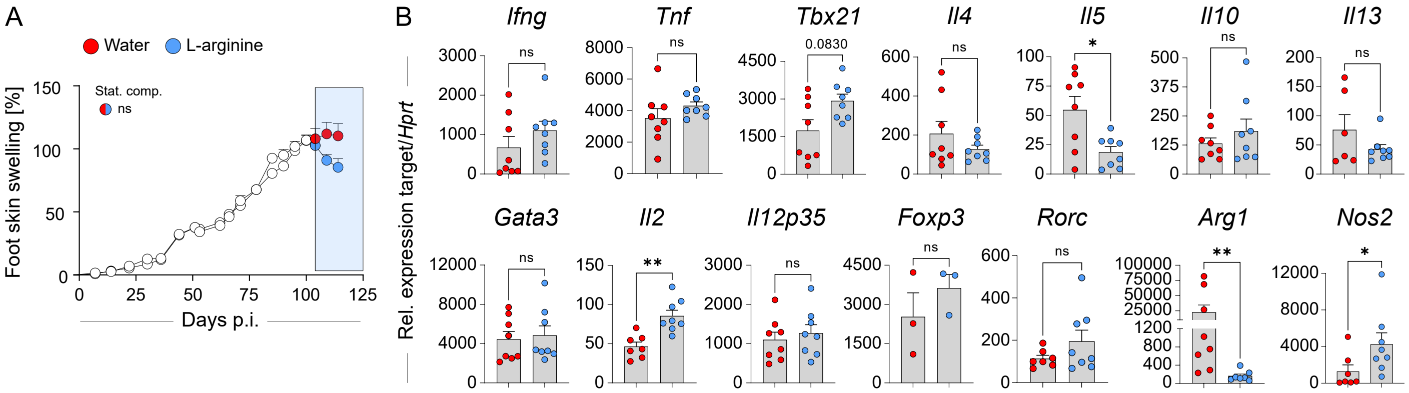


**Fig. S8: Th1 response is slightly enhanced in the dLN of L-arginine-treated mice. (A)** *L. mexicana*-infected C57BL/6N WT mice with chronic CL lesions (100% increase of skin swelling) were split into two groups that either received L-arginine-containing drinking water or remained on normal water (control). Analyses were performed at day 10-15 after initiation of L-arginine treatment (1 of 5 independent experiments). **(B)** mRNA expression (assessed by RT-qPCR) in dLN of *L. mexicana*-infected mice at day 10-15 of L-arginine treatment, which was started when the increase of skin lesion swelling had reached 100%. Data are presented as mean ± SEM (3–8 mice per group). Statistical significance was determined using a two-tailed Mann-Whitney U test **(B)**. Comparisons between respective mouse groups were performed using a mixed-model two-way repeated measures ANOVA. Statistical significance between groups is illustrated using half-circles with colours of the compared groups **(A)**. Significance levels are indicated according to the P value. Levels of significance: p > 0.05 (ns, not significant); *p < 0.05; **p < 0.01; ***p < 0.001; ****p < 0.0001.

**Table S1: Gene-specific assays for quantitative real-time PCR**

The following gene-specific assays (TaqMan Gene Expression Assays, Thermo Fisher Scientific) were used for quantitative real-time PCR:

<u>Mouse</u>

*Arg1* (Mm00475988_m1), *Arg1* exon 7-8 (Mm01190441_g1), *Arg2* (Mm00477592_m1), *Ccl2* (Mm00441242_m1), *Ccl3* (Mm99999057_m1), *Ccl5* (Mm01302427_m1), *Ccl7* (Mm00443113_m1), *Ccl8* (Mm01297183_m1), *Ccl11* (Mm00441238_m1), *Ccl24* (Mm00444701_m1), *Cxcl1* (Mm04207460_m1), *Cxcl2* (Mm00436450_m1), *Cxcl9* (Mm00434946_m1), *Cxcl10* (Mm00445235_m1), *Cxcl11* (Mm00444662_m1), *FoxP3* (Mm00475162_m1), *Gata3* (Mm00484683_m1), *Hprt-1* (Mm00446968_m1), *Ifng* (Mm00801778_m1), *Il2* (Mm00434256_m1), *Il4* (Mm00445259_m1), *Il5* *(*Mm00439646_m1), *Il10* (Mm00439616_m1), *Il12p35* (Mm00434165_m1), *Il13* (Mm00434204_m1), *Mhc2* (Mm00439216_m1), *Nos2* (Mm00440485_m1), Rorc (Mm01261022_m1), *Tnf* (Mm00443258_m1), *Tbx21* (Mm00450960_m1)

<u>Human</u>

*Arg1* (Hs00968979_m1), *Gapdh* (Hs02758991_g1), *Il4* (Hs00174122_m1), *Il10* (Hs00961622_m1), *Ifng* (Hs00989291_m1), *Nos2* (Hs01075529_m1), *Tnf* (Hs01113624_g1)

**Table S2: Antibodies used for Western Blot and ELISA**

***Western blot***

| **Primary antibody** | **Host species** | **Clone** | **Final dilution** | **Source** |
|---|---|---|---|---|
| NOS2 | Rabbit | Polyclonal | 1:20000 | Prof. C. Nathan, New York City, lot 3055D und lot 3055E |
| ARG1 | Goat | V20, Polyclonal | 1:500 | Santa Cruz Biotechnology, Dallas, TX, USA |
| ARG2 | Rabbit | H64, Polyclonal | 1:500 | Santa Cruz Biotechnology, Dallas, TX, USA |
| GRB2 | Mouse | Monoclonal | 1:500 | BD Transduction laboratories, BD Bioscieneces |
| **Secondary antibody** | **Host species** | **Clone** | **Final dilution** | **Company** |
| Rabbit POX (HRP conjugated) | Goat | Complete IgG | 1:40000 | Dianova Hamburg, Germany |
| Mouse POX (HRP conjugated) | Goat | Complete IgG | 1:40000 | Jackson ImmunoResearch, Ely, Cambridgeshire, UK |
| Goat POX (HRP conjugated) | Donkey | Complete IgG | 1:40000 | Jackson ImmunoResearch, Ely, Cambridgeshire, UK |

***Antibodies used for ELISA***

| **Antibody** | **Clone** | **Conjugation** | **Final dilution** | **Company** |
|---|---|---|---|---|
| IFN-γ Capture Antibody | R4-6A2 | | 1:500 | All from Biolegend, San Diego, CA, USA |
| Rat anti-mouse IFN-γ | XMG1.2 | Biotin | 1:500 | |
| Streptavidin | | HRP | 1:2000 | |
| IL4 Capture Antibody | 11B11 | | 1:1000 | |
| Rat anti-mouse IL4 | BVD6-24G2 | Biotin | 1:1000 | |
| IL10 Capture Antibody | JES5-2A5 | | 1:250 | |
| Anti-mouse IL10 | JES5-16E3 | Biotin | 1:500 | |

**Table S3: Antibodies used for flow cytometry and cell sorting**

***Antibodies used for flow cytometry***

| Target | Fluoro-chrome | Clone | Target species | Final dilution | Isotype | Company |
|---|---|---|---|---|---|---|
| ARG1 | APC | A1exF5 | mouse/ human | 1:50 | rat IgG2a, κ | Invitrogen |
| CD3 | PerCP-Cy5.5 | 17A2 | mouse | 1:100 | rat IgG2b, κ | BioLegend |
| CD3 | FITC | 145-2C11 | mouse | 1:100 | hamster IgG1, κ | eBioscience |
| CD4 | PE-Cy7 | GK1.5 | mouse | 1:100 | rat IgG2b, κ | eBioscience |
| CD4 | BV421 | GK1.5 | mouse | 1:100 | rat IgG2b, κ | BioLegend |
| CD11b | APC | M1/70 | mouse/ human | 1:200 | rat IgG2b, κ | BioLegend 101212 |
| CD11b | APC eF780 | M1/70 | mouse | 1:200 /1:100 | rat IgG2b, κ | eBioscience |
| CD11b | BUV395 | M1/70 | mouse | 1:200 | rat DA IgG2b, κ | BD |
| CD11b | BUV737 | M1/70 | mouse | 1:200 | rat IgG2b, κ | BD |
| CD16/ 32 | TruStain fcx | 93 | mouse | 1:100 | rat IgG2A, λ | BioLegend |
| IFNγ | APC | XMG1.2 | mouse | 1:100 | rat IgG1, κ | BioLegend |
| Ly6G | BV421 | 1A8 | mouse | 1:200 | rat IgG2a, κ | BioLegend |
| Ly6G | APC eF780 | 1A8 | mouse | 1:200/1:100 | rat IgG2a, κ | eBioscience |
| NK1.1 | BV421 | PK136 | mouse | 1:100 | mouse IgG2a, κ | BioLegend |
| NK1.1 | PE-Cy7 | PK136 | mouse | 1:100 | mouse IgG2a, κ | BioLegend 108741 |

| | | | | | | |
|---|---|---|---|---|---|---|
| NOS2 | PerCP-eF710 | CXNFT | mouse | 1:100 | rat IgG2a, κ | Invitrogen |
| NOS2 | PE | CXNFT | mouse | 1:100 | rat IgG2a, κ | eBioscience |
| SiglecF | PE-eF610 | 1RNM44N | mouse | 1:200 | rat IgG2a, κ | eBioscience |
| SiglecF | PE-Dazzle 594 | S17007L | mouse | 1:400 or 1:200 | rat IgG2a, κ | BioLegend |
| SiglecF | BV421 | E50-2440 | mouse | 1:100 | mouse IgG2a, κ | BD |
| Strep | BV785 | | | 1:100 | | Biolegend 405249 |
| TCRγδ | PE | GL3 | mouse | 1:100 | armenian hamster IgG | BioLegend |
| TCRγδ | Biotin | GL3 | mouse | 1:100 | armenian hamster IgG | BioLegend 118103 |

***Antibody used for ARG1$^+$NOS2$^+$ macrophage sorting***

| **Target** | **Fluoro chrome** | **Clone** | **Target species** | **Final dilution** | **Isotype** | **Company** |
|---|---|---|---|---|---|---|
| ARG1 | APC | A1exF5 | mouse/ human | 1:50 | rat IgG2a, κ | Invitrogen |
| NOS2 | PE | CXNFT | mouse | 1:100 | rat IgG2a, κ | eBioscience |
| CD11b | APC eF780 | M1/70 | mouse | 1:200 | rat IgG2b, κ | eBioscience |
| SiglecF | BV421 | E50-2440 | mouse | 1:100 | mouse IgG2a, κ | BD |

**Table S4: Antibodies used for CLSFM**

| **Primary antibody** | **Host species** | **Clone or specification** | **Final dilution** | **Company** |
|---|---|---|---|---|
| ARG1 | goat | V20, polyclonal | 1:100 | Santa Cruz Biotechnology, Dallas, TX, USA |
| NOS2 | rabbit | iNOS16 R1949, serum, polyclonal | 1:2000 | Prof. C. Nathan, New York City, lot 3055D und lot 3055E |
| *L. mexicana* | human | polycloncal serum D.P. (20.09.2013) | 1:400 | Serum from patient infected with *L. mexicana* |
| Nitro-tyrosine | rabbit | polyclonal serum | 1:1000 | Sigma Aldrich St. Louis, MO, USA |
| **Secondary antibody** | **Host species** | **Clone or specification** | **Final dilution** | **Company** |
| anti-goat A488 | donkey | H+L(ab)2 Fragment | 1:100 | Dianova Hamburg, Germany |
| anti-rabbit Rhodamine Red X | donkey | H+L(ab)2 Fragment | 1:100 | Dianova Hamburg, Germany |
| anti-human A647 | donkey | H+L(ab)2 Fragment | 1:100 | Dianova Hamburg, Germany |
| Anti-rabbit A488 | goat | H+L (Complete IgG) | 1:1000 | Invitrogen, Carlsbad, CA, USA |

**Table S5: Metabolomic analyses by LC/MS**

*m/z transitions of all compounds measured in tissue lysates*

*m/z transitions of all compounds measured in tissue lysates*

| **Analyte** | **Precursor (m/z)** | **Daughter ion (m/z)** | **Internal standard** |
|---|---|---|---|
| Agmatine | 266.1 | 173.1 | d8-Agmatine |
| Agmatine Q | 266.101 | 207.1 | d8-Agmatine 2 |
| Alanine | 225.2 | 44.2 | d4-Alanine |
| alpha-AAA | 297.1 | 144.2 | d4-Alanine |
| Arginine | 310.1 | 217 | d7-Arginine |
| Aspartate | 269.2 | 116.2 | d3-Aspartic acid |
| Citrulline | 311.2 | 113.1 | d2-Citrulline |
| Citrulline | 311.21 | 113.1 | d2-Citrulline |
| Glutamine | 282.2 | 130 | d5-Glutamine |
| Glutamic acid | 283.2 | 130.2 | d5-Glutamic acid |
| Glycine | 211.21 | 75.9 | 13C2-15N-Glycine |
| Histidine | 291.11 | 110.2 | 13C6-Histidine |
| Isoleucine | 267.3 | 69 | 13C615N-Isoleucine |
| Leucine | 267.3 | 43 | d3-Leucine |
| Lysine | 417.21 | 324.2 | 13C6-Lysine |
| Methionine | 285.1 | 104.2 | d3-Methionine |
| OH-Proline | 267.1 | 68 | d7-Proline |
| Ornithine | 403.21 | 310.2 | d6-Ornithine |

| | | | |
|---|---|---|---|
| Phenylalanine | 301.21 | 120.2 | d5-Phenylalanine |
| Proline | 251.21 | 70.3 | d7-Proline |
| Putrescine | 266.1 | 113.9 | d8-Putrescine |
| Serine | 241.2 | 60 | d3-Serine |
| Spermidine | 551.2 | 193.2 | d6-Spermidine |
| Spermine | 608.3 | 193.2 | d20 Spermine Q |
| Spermine Q | 473.3 | 193.2 | d20-Spermine |
| Threonin | 255.2 | 74.1 | 13C4-15N-Threonine |
| Tryptophan | 340.21 | 188.2 | d8-Tryptophan |
| Tyrosine | 317.21 | 136.1 | d4-Tyrosine |
| Valine | 253.21 | 72.2 | d8-Valine |

*m/z transitions of all compounds measured in plasma*

| **m/z Q1** | **m/z Q3** | **Name** | **DP [V]** | **EP [V]** | **CE [eV]** | **CXP [V]** |
|---|---|---|---|---|---|---|
| 175,9 | 71 | Arginin | 40 | 10 | 28 | 11 |
| 175,9 | 60 | Arginin_Q1 | 40 | 10 | 18 | 11 |
| 180,9 | 74 | Arginin_ISTD | 40 | 10 | 28 | 11 |
